# Evolutionary-scale prediction of atomic level protein structure with a language model

**DOI:** 10.1101/2022.07.20.500902

**Authors:** Zeming Lin, Halil Akin, Roshan Rao, Brian Hie, Zhongkai Zhu, Wenting Lu, Nikita Smetanin, Robert Verkuil, Ori Kabeli, Yaniv Shmueli, Allan dos Santos Costa, Maryam Fazel-Zarandi, Tom Sercu, Salvatore Candido, Alexander Rives

## Abstract

Artificial intelligence has the potential to open insight into the structure of proteins at the scale of evolution. It has only recently been possible to extend protein structure prediction to two hundred million cataloged proteins. Characterizing the structures of the exponentially growing billions of protein sequences revealed by large scale gene sequencing experiments would necessitate a break-through in the speed of folding. Here we show that direct inference of structure from primary sequence using a large language model enables an order of magnitude speed-up in high resolution structure prediction. Leveraging the insight that language models learn evolutionary patterns across millions of sequences, we train models up to 15B parameters, the largest language model of proteins to date. As the language models are scaled they learn information that enables prediction of the three-dimensional structure of a protein at the resolution of individual atoms. This results in prediction that is up to 60x faster than state-of-the-art while maintaining resolution and accuracy. Building on this, we present the ESM Metagenomic Atlas. This is the first large-scale structural characterization of metagenomic proteins, with more than 617 million structures. The atlas reveals more than 225 million high confidence predictions, including millions whose structures are novel in comparison with experimentally determined structures, giving an unprecedented view into the vast breadth and diversity of the structures of some of the least understood proteins on earth.

## 1. Introduction

The sequences of proteins at the scale of evolution contain an image of biological structure and function. This is because the biological properties of a protein act as constraints on the mutations to its sequence that are selected through evolution, recording structure and function into evolutionary patterns (1–3). Within a protein family, structure and function can be inferred from the patterns in sequences (4, 5). This insight has been central to progress in computational structure prediction starting from classical methods (6, 7), through the introduction of deep learning (8–11), up to the present state-of-the-art (12, 13).

The idea that biological structure and function are reflected in the patterns of protein sequences has also motivated a new line of research on evolutionary scale language models (14). Beginning with Shannon’s model for the entropy of text (15), language models of increasing complexity have been developed to fit the statistics of text, culminating in modern large-scale attention based architectures (16–18). Language models trained on the amino acid sequences of millions of diverse proteins have the potential to learn patterns across all of them. This idea contrasts with the standard basis for inference from protein sequences, which begins from a multiple sequence alignment summarizing the evolutionary patterns in related proteins.

In artificial intelligence, language models of text, despite the simplicity of their training objectives, such as filling in missing words or predicting the next word, are shown to develop emergent capabilities that are connected to the underlying meaning of the text. These capabilities develop as a function of scale, with greater capabilities emerging as computation, data, and number of parameters increase. Modern language models containing tens to hundreds of billions of parameters develop abilities such as few-shot language translation, commonsense reasoning, and mathematical problem solving, all without explicit supervision (19–22). These observations raise the possibility that a parallel form of emergence might be exhibited by language models trained on protein sequences.

We posit that the task of filling in missing amino acids in protein sequences across evolution will require a language model to learn something about the underlying structure that creates the patterns in the sequences. As the representational capacity of the language model and the diversity of protein sequences seen in its training increase, we expect that deep information about the biological properties of the protein sequences could emerge, since those properties give rise to the patterns that are observed in the sequences. To study this kind of emergence we scale language models from 8 million parameters up to 15 billion parameters. We discover that atomic resolution structure prediction emerges and continues to improve in language models over the four orders of magnitude in parameter scale. Strong correlations between the language model’s understanding of the protein sequence (perplexity) and the accuracy of the structure prediction reveal a close link between language modeling and the learning of structure.

We show that language models enable fast end-to-end atomic resolution structure prediction directly from sequence. Our new approach leverages the evolutionary patterns captured by the language model to produce accurate atomic level predictions. This removes costly aspects of current state-of-the-art structure prediction pipelines, eliminating the need for a multiple sequence alignment, while at the same time greatly simplifying the neural architecture used for inference. This results in an improvement in speed of up to 60x on the inference forward pass alone, while also removing the search process for related proteins entirely, which can take over 10 minutes with the high-sensitivity pipelines used by AlphaFold (12) and RosettaFold (13), and which is a significant part of the computational cost even with new lower sensitivity fast pipelines (23). In practice this means the speedup over the state-of-the-art prediction pipelines that are in use is up to one to two orders of magnitude.

This makes it possible to expand structure prediction to metagenomic proteins. The last decade has seen efforts to expand knowledge of protein sequences to the immense microbial natural diversity of the earth through metagenomic sampling. These efforts have contributed to an exponential growth in the size of protein sequence databases, which now contain billions of proteins (24–26). While computational structural characterizations have recently been completed for ∼20K proteins in the human proteome (27), and the ∼200M cataloged proteins of Uniprot (28), the vast scale of metagenomic proteins represents a far greater challenge for structural characterization. The extent and diversity of metagenomic structures is unknown and is a frontier for biological knowledge, and a potential source of new discoveries for medicine and biotechnology (29–31).

We present the first evolutionary scale structural characterization of a metagenomic resource, folding practically all sequences in MGnify90 (25), over 617M proteins. We are able to complete this characterization in 2 weeks on a heterogeneous cluster of 2,000 GPUs, demonstrating scalability to far larger databases. High confidence predictions are made for over 225M structures, revealing and characterizing regions of metagenomic space distant from existing knowledge with the vast majority (76.8%) of high confidence predictions being separate from UniRef90 (32) by at least 90% sequence identity, and tens of millions of predictions (12.6%) without a match to experimentally determined structures. These results give the first large-scale view into the vast extent and diversity of metagenomic protein structures.

All predictions can be accessed in the ESM Metage-nomic Atlas (https://esmatlas.com) open science resource.

## 2. Atomic resolution structure emerges in language models trained on protein sequences

We begin with a study of the emergence of high resolution protein structure. We train a new family of transformer protein language models, ESM-2, at scales from 8 million parameters up to 15 billion parameters. Relative to our previous generation model ESM-1b, ESM-2 introduces improvements in architecture, training parameters, and increases computational resources and data (Appendices A.1.1 and A.2). The resulting ESM-2 model family significantly outperforms previously state-of-the-art ESM-1b (a ∼650 million parameter model) at a comparable number of parameters, and on structure prediction benchmarks it also outperforms other recent protein language models (Table S1).

The ESM-2 language models are trained with the masked language modeling objective (18), which trains the model to predict the identity of randomly selected amino acids in a protein sequence by observing their context in the rest of the sequence. This causes the model to learn dependencies between the amino acids. Although the training objective itself is simple and unsupervised, performing well on this task over millions of evolutionarily diverse protein sequences requires the model to internalize sequence patterns across evolution. We expect that this training will also cause structure to materialize since it is linked to the sequence patterns. ESM-2 is trained over sequences in the UniRef (32) protein sequence database. During training, sequences are sampled with even weighting across ∼43 million UniRef50 training clusters from ∼138 million UniRef90 sequences so that over the course of training the model sees ∼65 million unique sequences.

As we increase the scale of ESM-2 from 8 million to 15 billion parameters, we observe large improvements in the fidelity of its modeling of protein sequences. This fidelity can be measured using perplexity, which ranges from 1 for a perfect model to 20 for a model that makes predictions at random. Intuitively, the perplexity describes the number of amino acids the model is choosing between for each prediction. Fig. S1 shows perplexity for the ESM-2 family as a function of the number of training updates, evaluated on a set of ∼500K UniRef50 clusters that have been held out from training. Comparisons are performed at 270k training steps for all models in this section. The fidelity continues to improve as the parameters increase up to the largest model. The 8M parameter model has a perplexity of 10.45, and the 15B model reaches a perplexity of 6.37, indicating a large improvement in the understanding of protein sequences with scale.

This training also results in the emergence of structure in the models. Since ESM-2’s training is only on sequences, any information about structure that develops must be the result of representing the patterns in sequences. Transformer models trained with masked language modeling, are known to develop attention patterns that correspond to the residue-residue contact map of the protein (33, 34). We examine how this low resolution picture of protein structure emerges as a function of scale. We use a linear projection to extract the contact map from the attention patterns of the language model (Appendix A.2.1). The precision of the top L (length of the protein) predicted contacts (long range contact precision) measures the correspondence of the attention pattern with the structure of the protein. Attention patterns develop in ESM-2 that correspond to tertiary structure (Fig. 1A), and scaling leads to large improvements in the understanding of structure (Fig. 1B). The accuracy of the predicted contacts varies as a function of the number of evolutionarily related sequences in the training set. Proteins with more related sequences in the training set have steeper learning trajectories with respect to model scale (Fig. 1C). This means that improvement on sequences with high evolutionary depth saturates at lower model scales, and improvement on sequences with low evolutionary depth continues as models increase in size.

**Figure 1.**
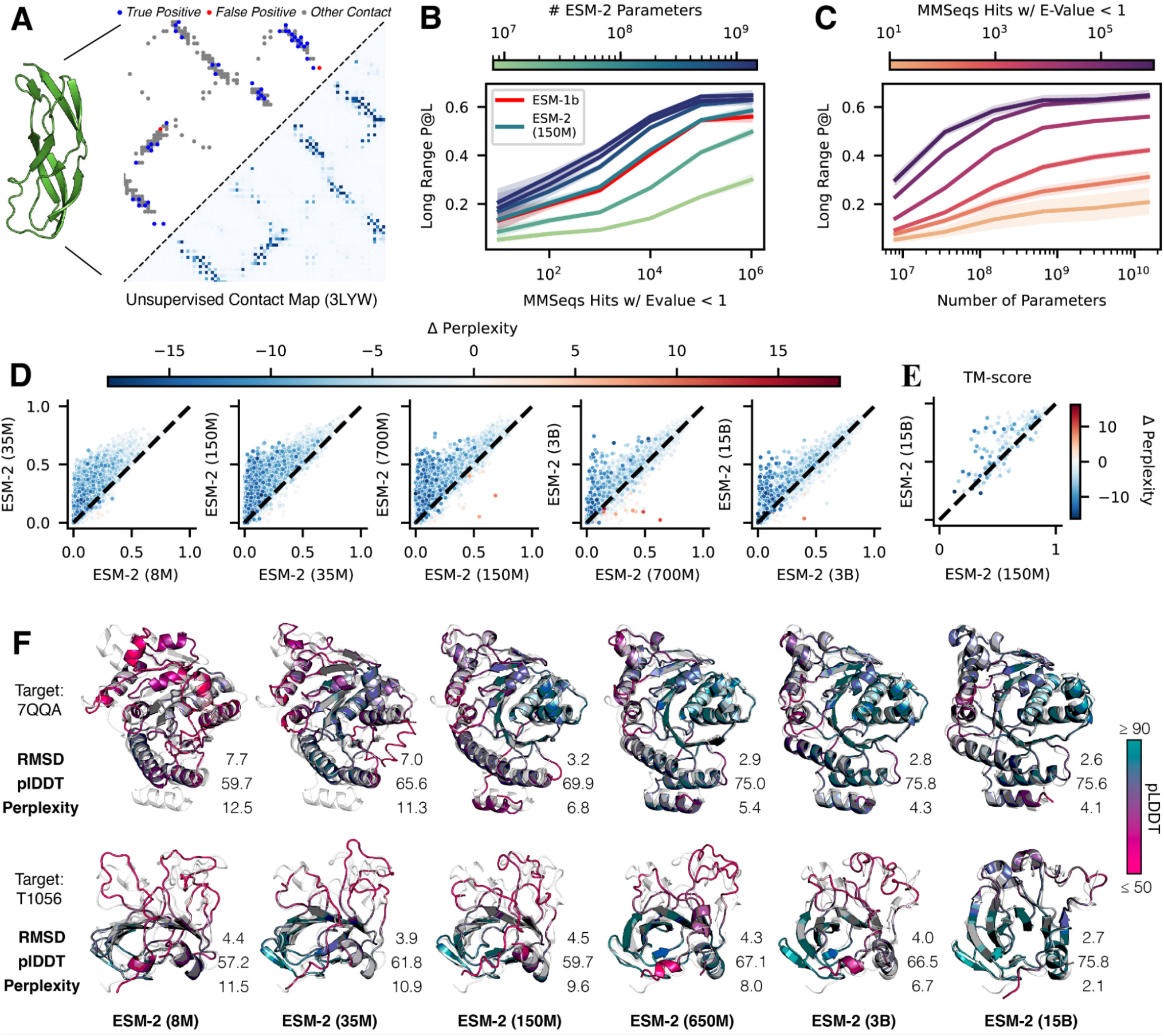
Emergence of structure when scaling language models to 15 billion parameters. (A) Predicted contact probabilities (bottom right) and actual contact precision (top left) for 3LYW. A contact is a positive prediction if it is within the top-L most likely contacts for a sequence of length L. (B, C, D) Unsupervised contact prediction (long range precision at L) for all scales of the ESM-2 model. (B) Performance binned by the number of MMseqs hits when searching the training set. Larger ESM-2 models perform better at all levels; the 150M parameter ESM-2 model is comparable to the 650M parameter ESM-1b model. (C) Trajectory of improvement as model scale increases for sequences with different numbers of MMseqs hits. (D) Left-to-right shows models from 8M to 15B parameters, comparing the smaller model (x-axis) against the next larger model (y-axis) via unsupervised contact precision. Points are PDB proteins colored by change in pseudo-perplexity for the sequence between the smaller and larger model. Sequences with large changes in contact prediction performance also exhibit large changes in language model understanding measured by pseudo-perplexity. (E) TM-score on combined CASP14 and CAMEO test sets. Predictions are made using structure module-only head on top of language models. Points are colored by the change in pseudo-perplexity between the models. (F) Structure predictions on CAMEO structure 7QQA and CASP target 1056 at all ESM-2 model scales, colored by pLDDT (pink = low, teal = high). For 7QQA, prediction accuracy improves at the 150M parameter threshold. For T1056, prediction accuracy improves at the 15B parameter threshold.

For individual proteins, we often observe non-linear improvements in the accuracy of the contact prediction as a function of scale. Fig. 1D plots the change in the distribution of long range contact precision at each transition to a higher level of scale. At each step there is an overall shift in the distribution toward better performance. Also at each transition, there is a subset of proteins that undergo significant improvement. In Fig. 1D these are in the upper left of each plot, far from the diagonal. The accuracy of the contact map prediction and perplexity are linked, with proteins undergoing large changes in contact map accuracy also undergoing large changes in perplexity (NDCG = 0.87, Appendix A.2.6). This link indicates that the language modeling objective is directly correlated with the materialization of the folded structure in the attention maps.

We investigate whether high resolution structure at an atomic level also develops. To identify atomic resolution information in the model, we project out spatial coordinates for each of the atoms from the internal representations of the language model using an equivariant transformer (Appendix A.3.3). This projection is fit using experimentally determined protein structures from PDB (35), and evaluated on 194 CAMEO proteins (36) and 51 CASP14 proteins (37). TM-score, which ranges from 0 to 1, measures the accuracy of the projection in comparison to the ground truth structure, with a value of 0.5 corresponding to the threshold for correctly predicting the fold (38). The evaluation uses a temporal cutoff, ensuring that the proteins used for testing are held out from those used in fitting the projection. This makes it possible to measure how atomic level information emerges in the representations as a function of the parameter scale.

We discover that an atomic resolution structure prediction can be projected from the representations of the ESM-2 language models. The accuracy of this projection improves with the scale of the language model. The 15 billion parameter model reaches a TM-score of 0.72 on the CAMEO test set and 0.55 on the CASP14 test set, a gain of 14% and 17% respectively relative to the the 150 million parameter ESM-2 model (Fig. 1E). At each increase in scale a subset of proteins undergo large changes in accuracy. For example, the protein 7QQA improves in RMSD from 7.0 to 3.2 when scale is increased from 35M to 150M parameters, and the CASP target T1056 improves in RMSD from 4.0 to 2.6 when scale is increased from 3B to 15B parameters (Fig. 1F). Before and after these jumps, changes in RMSD are much smaller. Across all models (Table S1) there is a correlation of -0.99 between validation perplexity and CASP14 TM-score, and -1.00 between validation perplexity and CAMEO TM-score indicating a strong connection between the understanding of the sequence measured by perplexity and the atomic resolution structure prediction. Additionally there are strong correlations between the low resolution picture of the structure that can be extracted from the attention maps and the atomic resolution prediction (0.96 between long range contact precision and CASP14 TM-score, and 0.99 between long range contact precision and CAMEO TM-score). These findings connect improvements in language modeling with the increases in low resolution (contact map) and high resolution (atomic level) structural information.

## 3. Accelerating accurate atomic resolution structure prediction with a language model

Language models greatly accelerate state-of-the-art high resolution structure prediction. The language model internalizes evolutionary patterns linked to structure, eliminating the need for external evolutionary databases, multiple sequence alignments, and templates. We find that the ESM-2 language model generates state-of-the-art three-dimensional structure predictions directly from the primary protein sequence. This results in a speed improvement for structure prediction of more than an order of magnitude while maintaining high resolution accuracy.

We develop ESMFold, a fully end-to-end single sequence structure predictor, by training a folding head for ESM-2 (Fig. 2A). At prediction time the sequence of a protein is input to ESM-2. The sequence is processed through the feedforward layers of the language model, and the model’s internal states (representations) are passed to the folding head. The head begins with a series of folding blocks. Each folding block alternates between updating a sequence representation and a pairwise representation. The output of these blocks is passed to an equivariant transformer structure module, and three steps of recycling are performed before outputting a final atomic-level structure and predicted confidences (Appendix A.3.1). This architecture represents a major simplification in comparison to current state-of-the-art structure prediction models which deeply integrate the multiple sequence alignment into the neural network architecture through an attention mechanism operating across the rows and columns of the MSA (12, 40).

**Figure 2.**
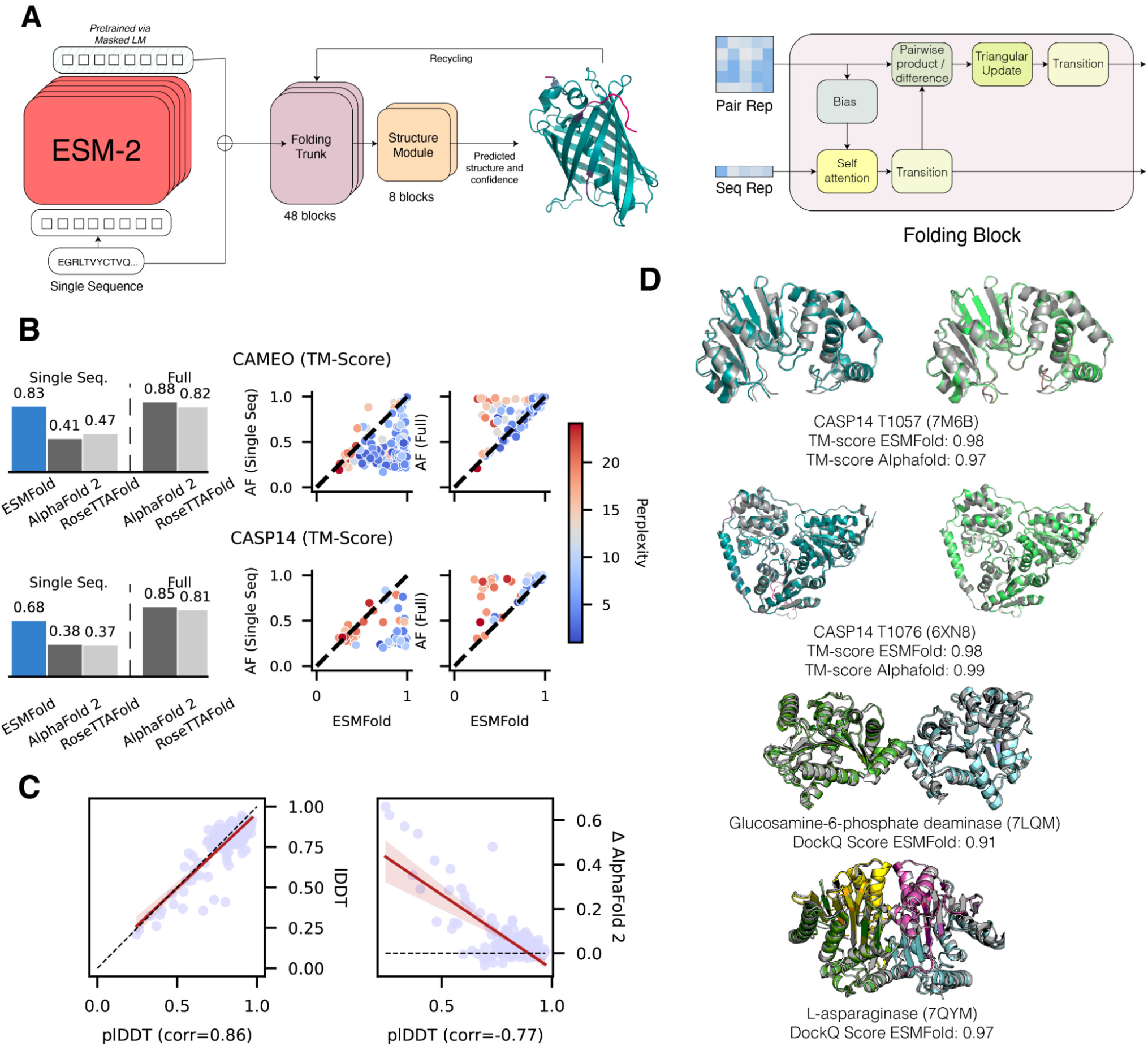
Single sequence structure prediction with ESMFold. (A) ESMFold model architecture. Arrows show the information flow in the network from the language model to the folding trunk to the structure module which outputs 3D coordinates and confidences. (B) ESMFold produces accurate atomic resolution predictions, with similar accuracy to RosettaFold on CAMEO. When MSAs are ablated for AlphaFold and RosettaFold, performance of the models degrades. Scatter-plots compare ESMFold (x-axis) predictions with AlphaFold2 (y-axis), colored by language model perplexity. Proteins with low perplexity score similarly to AlphaFold2. (C) Model pLDDT vs. true LDDT (left) and relative performance against AlphaFold (right) on CAMEO. pLDDT is a well calibrated estimate of prediction accuracy. (D) Top shows test-set predictions of ESMFold in teal, ground truth in gray, and AlphaFold2 predictions in green. Pink shows low predicted LDDT for both ESMFold and AlphaFold2. Bottom shows complex predictions on a dimer (7LQM) and a tetramer (7QYM); ESMFold predictions are colored by chain ID and overlaid on ground truth (gray). DockQ (39) scores are reported for the interactions; in the case of the tetramer 7QYM, the score is the average of scores over interacting chain-pairs.

Our approach results in a significant improvement in prediction speed. On a single NVIDIA V100 GPU, ESMFold makes a prediction on a protein with 384 residues in 14.2 seconds, 6x faster than a single AlphaFold2 model. On shorter sequences the improvement increases up to ∼60x (Fig. S2). The search process for related sequences, required to construct the MSA, can take over 10 minutes with the high sensitivity protocols used by the published versions of AlphaFold and RosettaFold; this can be reduced to less than 1 minute, although with reduced sensitivity (23).

We train the folding head on ∼25K clusters covering a total of ∼325K experimentally determined structures from the PDB, further augmented with a dataset of ∼12M structures we predicted with AlphaFold2 (Appendix A.1.2). The model is trained with the same losses that are used for AlphaFold (41). To evaluate the accuracy of structure predictions we use test sets that are held out from the training data by a May 2020 cutoff date; as a result all structures that are used in evaluation are held out from the training, and the evaluation is representative of the performance that would be expected in regular usage as a predictive model on the kinds of structures that are selected by experimentalists for characterization. This also makes it possible to compare with AlphaFold and RosettaFold since these models also have not been trained on structures deposited after May 2020. We use two test sets: the CAMEO test set consists of 194 structures used in the ongoing CAMEO assessment (between April 2022 to June 2022); the CASP14 test set consists of 51 publicly released structures that have been selected for their difficulty for the biannual structure prediction competition.

We compare results on these evaluation sets to AlphaFold2 and RosettaFold (Fig. 2B). ESMFold achieves an average TM-score of 0.83 on CAMEO and 0.68 on CASP14. Using the search protocols released with AlphaFold2, including MSAs and templates, AlphaFold2 achieves 0.88 and 0.85 on CAMEO and CASP14 respectively. ESMFold achieves competitive accuracy with RosettaFold on CAMEO, which averages a TM-score of 0.82. When evaluating AlphaFold2 and RosettaFold on single sequences by ablating the multiple sequence alignment, performance degrades substantially, and falls well below that of ESMFold. Note that this is an artificial setting as AlphaFold2 has not been explicitly trained for single sequences, however it has recently emerged as important in protein design, where these models have been used with single sequence inputs for *de novo* protein design (42–44).

Because the language model is the critical component of ESMFold, we test how well differences in the language model’s understanding of a sequence correspond to changes in the accuracy of structure prediction. The performance of ESMFold on both test sets is well correlated with the perplexity of the language model. On the CAMEO test set, language model perplexity has a Pearson correlation of -0.55 with the TM-score between the predicted and experimental structures; on CASP14, the correlation is -0.67 (Fig. 2B). The relationship between perplexity and structure prediction suggests that improving the language model is key to improving single-sequence structure prediction accuracy, consistent with observations from the scaling analysis (Figs. 1D and 1E). Additionally, this means the language model’s perplexity for a sequence can be used to predict the quality of the ESMFold structure prediction.

Ablation studies indicate that the language model representations are critical to ESMFold performance (Fig. S3). With a folding trunk of 8 blocks, performance on the CAMEO test set is 0.74 LDDT (baseline). Without the language model, this degrades substantially, to 0.58 LDDT. When removing the folding trunk entirely (i.e. only using the language model and the structure module), performance degrades to 0.66 LDDT. Other ablations: only 1 block of a structure module, turning off recycling, not using AlphaFold2 predicted structures as distillation targets, or not using triangular updates, result in small performance degradations (change in LDDT of -0.01 to -0.04).

ESMFold provides state-of-the-art structure prediction accuracy, matching AlphaFold2 performance (*<* 0.05 LDDT difference) on more than half the proteins (Fig. 2B). We find that this is true even on some large proteins—T1076 is an example with 0.98 TM-score and 540 residues (Fig. 2D). Parts of structure with low accuracy do not differ significantly between ESMFold and AlphaFold, suggesting that language models are learning information similar to that contained in MSAs. We also observe that ESMFold is able to make good predictions for components of homo- and heterodimeric protein-protein complexes (Fig. 2D). In a comparison with AlphaFold-Multimer (45) on a dataset of 2,978 recent multimeric complexes deposited in the PDB, ESMFold achieves the same qualitative DockQ (39) categorization for 53.2% of chain pairs, despite not being trained on protein complexes (Fig. S4).

Confidence is well calibrated with accuracy. ESMFold reports confidence in the form of predicted-LDDT. This confidence correlates well with the accuracy of the prediction, and for high-confidence predictions (pLDDT *>* 0.7) accuracy is comparable to AlphaFold2 (ESMFold LDDT=0.83, AlphaFold2 LDDT=0.85 on CAMEO) (Figs. 2C and S5). High-confidence predictions approach experimental-level accuracy. On the CAMEO test set, ESMFold predictions have a median all-atom RMSD_95_ (root-mean-squared deviation at 95% residue coverage) of 1.91Å and backbone RMSD_95_ of 1.33Å. When confidence is very high (pLDDT > 0.9), predictions have median all-atom RMSD_95_ of 1.42Å and backbone RMSD_95_ of 0.94Å. This means the confidence can be used to predict how likely it is that a given structure prediction will match the true structure if it were to be experimentally determined.

## 4. Evolutionary-scale structural characterization of metagenomics

This fast and high resolution structure prediction capability enables the first full-scale structural characterization of a large metagenomic sequence resource. We fold over 617 million sequences from the MGnify90 database (25). This is the entirety of the sequences of length 20 to 1024, and covers 99% of all the sequences in MGnify90. Overall, this large-scale characterization produces ∼365 million predictions with good confidence (mean pLDDT *>* 0.5 and pTM *>* 0.5) corresponding to ∼59% of the database, and ∼225 million predictions with high confidence (mean pLDDT > 0.7 and pTM *>* 0.7) corresponding to ∼36% of total structures folded (Fig. 3). We were able to complete the predictions in 2 weeks on a cluster of approximately 2,000 GPUs (Appendix A.4.1).

**Figure 3.**
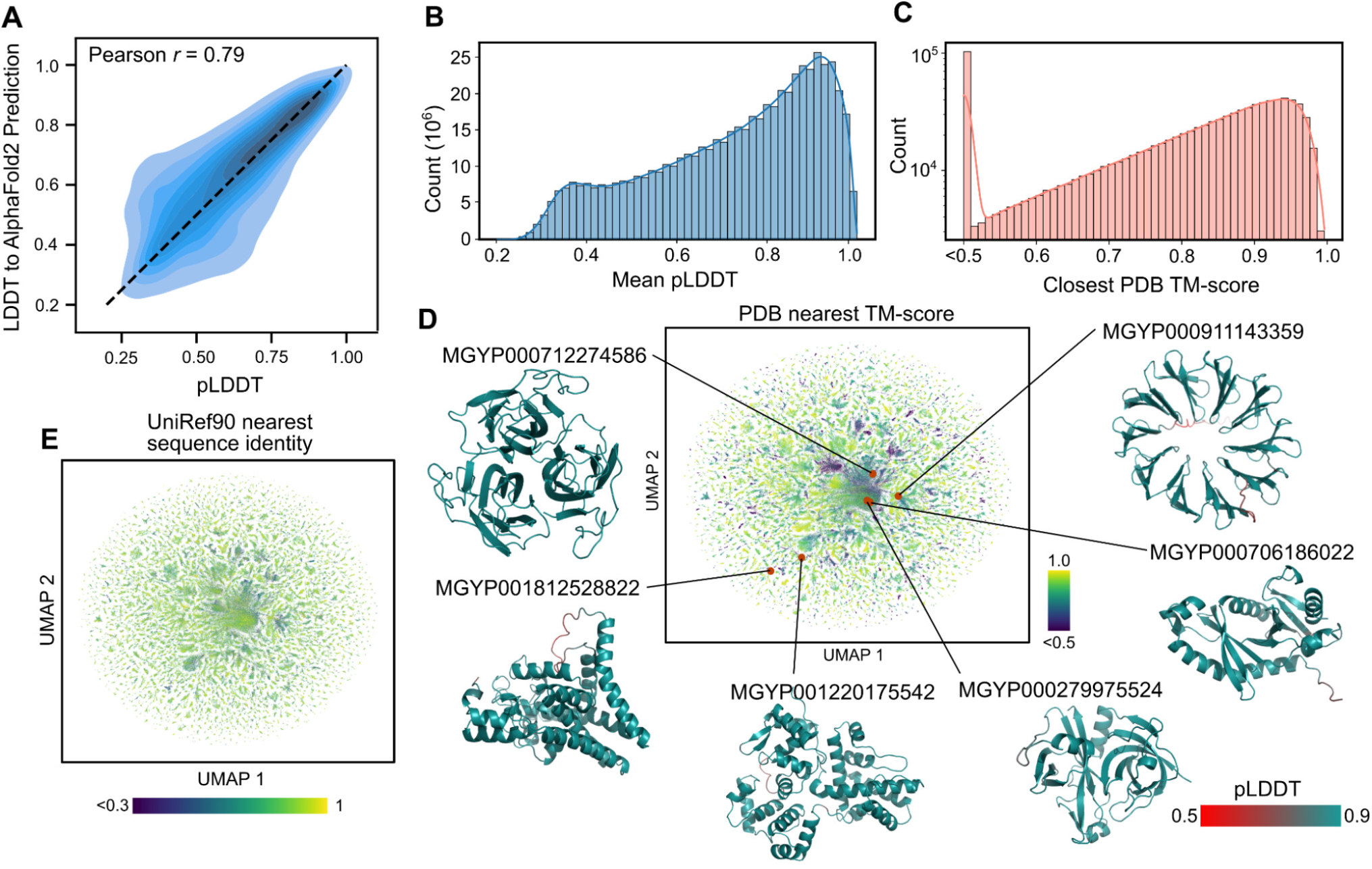
Mapping metagenomic structural space. (A) ESMFold calibration with AlphaFold2 for metagenomic sequences. Mean pLDDT is shown on the x-axis, and LDDT to the corresponding AlphaFold2 prediction is shown on the y-axis. Distribution is shown as a density estimate across a subsample of ∼4K sequences from the MGnify database. (B) The distribution of mean pLDDT values computed for each of ∼617 million ESMFold-predicted structures from the MGnify database. (C) The distribution of the TM-score to the most similar PDB structure for each of 1 million randomly sampled high confidence (mean pLDDT *>* 0.7 and pTM *>* 0.7) structures. Values were obtained by a Foldseek search (46). (D) This sample of 1 million high-confidence protein structures is visualized in two dimensions using the UMAP algorithm and colored according to distance from nearest PDB structure, where regions with low similarity to known structures are colored in dark blue. Example protein structures and their locations within the sequence landscape are provided; see also Fig. 4 and Table S2. (E) Additional UMAP plot in which the 1 million sequences are plotted according to the same coordinates as in (D) but colored by the sequence identity to the most similar entry in UniRef90 according to a blastp search.

For structure prediction at scale, it will be critical to distinguish well predicted proteins from those that are poorly predicted. In the previous section, we evaluated calibration against experimentally determined structures on held out test sets, finding that the model confidence is predictive of the agreement with experimentally determined structures. We also assess calibration against AlphaFold predictions on metagenomic proteins. On a random subset of ∼4K metagenomic sequences, there is a high correlation (Pearson r = 0.79) between ESMFold pLDDT and the LDDT to AlphaFold2 predictions (Fig. 3A). Combined with results on CAMEO showing that when confidence is very high (pLDDT *>* 0.9), ESMFold predictions often approach experimental accuracy, these findings mean that ESMFold’s confidence scores provide a good indication of the agreement with experimental structures and with the predictions that can be obtained from AlphaFold2. Across the ∼617 million predicted structures, ∼113 million structures meet the very high confidence threshold.

Many of our metagenomic structure predictions have high confidence (Fig. 3B) as well as a high degree of novelty (Figs. 3C to 3E). On a random sample of 1 million high confidence structures, 76.8% (767,580) of the proteins have a sequence identity below 90% to any sequence in UniRef90, indicating that these proteins are distinct from existing UniRef90 clusters (Fig. 3D). For 3.4% (33,521 proteins), no significant match is found in UniRef90 at all (Appendix A.4.2). Many structures are novel in comparison with experimentally determined structures. For 12.6% of the structures (125,765 proteins), no structure is found with TM-score over 0.5 in the PDB database (Figs. 3C and 3E), indicating that no experimentally determined structure with a similar fold could be identified. Relaxing this threshold to a TM-score of 0.7, reveals 25.4% (253,905 proteins) without similar structures in the PDB. For 2.6% (25,664) there is both low structural similarity (TM-score ≤ 0.5) and no close sequence homolog (*<* 30% identity) (Fig. 4A and Table S2). These results indicate that ESMFold effectively characterizes regions of the protein landscape that are distant from existing knowledge.

**Figure 4.**
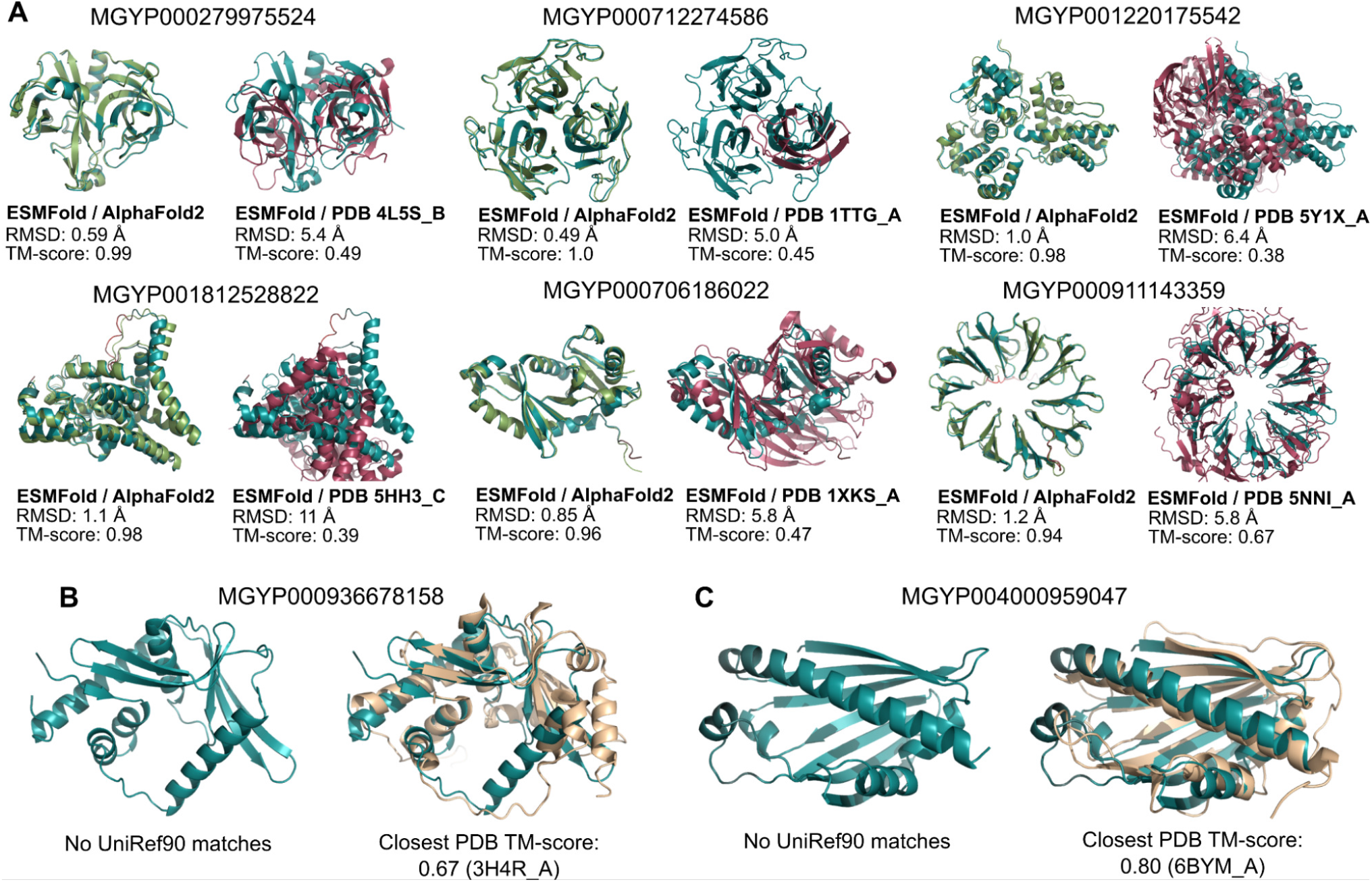
Example ESMFold structure predictions of metagenomic sequences. (A) Example predicted structures from six different metagenomic sequences; also see Table S2. Left of each subfigure: The prediction is displayed with the AlphaFold2 prediction (light green). Right of each subfigure: The prediction is displayed with the Foldseek-determined nearest PDB structure according to TM-score. (B, C) Examples of two ESMFold-predicted structures that have good agreement with experimental structures in the PDB but that have low sequence identity to any sequence in UniRef90. (B) The predicted structure of MGYP000936678158 aligns to an experimental structure from a bacterial nuclease (light brown, PDB: 3H4R), while (C) the predicted structure of MGYP004000959047 aligns to an experimental structure from a bacterial sterol binding domain (light brown, PDB: 6BYM).

Large scale structural characterization also makes it possible to identify structural similarities in the absence of sequence similarity. Many high-confidence structures with low similarity to UniRef90 sequences do have similar structures in the PDB. This remote homology often extends beyond the limit detectable by sequence similarity. For example, MGnify sequence MGYP000936678158 has no significant sequence matches to any entry in UniRef90, nor any significant matches via a jackhmmer (47) reference proteome search, but has a predicted structure conserved across many nucleases (PDB 5YET B, TM-score 0.68; PDB 3HR4 A, TM-score 0.67) (Fig. 4B and Table S2); similarly, MGnify sequence MGYP004000959047 has no significant UniRef90 or jackhmmer reference proteome matches but its predicted structure has high similarity to experimental structures of lipid binding domains (PDB 6BYM A, TM-score 0.80; PDB 5YQP B, TM-score 0.78) (Fig. 4C and Table S2). The ability to detect remote similarities in structure enables insight into function that cannot be obtained from the sequence.

All predicted structures are available in the ESM Metage-nomic Atlas (https://esmatlas.com) as an open science resource. Structures are available for bulk download, via a programmatic API, and through a web resource which provides search by sequence and by structure (46, 48). These tools facilitate both large scale and focused analysis of the full scope of the hundreds of millions of predicted structures.

## 5. Background

In this section, we provide a brief review of evolutionary scale language models. In Rives et al. (14) we found evidence that biological structure and function emerge in language models trained on protein sequences at the scale of evolution. Concurrently Bepler and Berger (49), Alley et al. (50), Heinzinger et al. (51) investigated LSTMs at a smaller scale and also found some evidence of biological properties in representations. Early models did not match performance of even simple evolutionary features on many tasks (52). Analysis of state-of-the-art evolutionary scale models such as ESM-1b and ProtTrans showed that low resolution structure, i.e., secondary structure (14, 53), and contact maps (14, 33, 34) could be recovered from representations.

Evolutionary scale models are also shown to perform unsupervised prediction of mutational effects (54, 55), and have recently been used in state-of-the-art applications, for example to predict the path of viral evolution (56, 57), and the clinical significance of gene variants (58). Several large scale models are now available as open source (14, 53, 59). Language models have been studied for end-to-end single sequence prediction of backbone structure (60).

## 6. Conclusions

Fast and accurate computational structure prediction has the potential to accelerate progress toward an era where it is possible to understand the structure of all proteins discovered in gene sequencing experiments. This promises new insights into the vast natural diversity of proteins, most of which is being newly discovered in metagenomic sequencing. To this end we have completed the first large-scale structural characterization of metagenomic proteins. This characterization reveals the structures of hundreds of millions proteins that have been previously unknown, millions of which are novel in comparison to experimentally determined structures.

As structure prediction continues to scale to larger numbers of proteins, the calibration of the model will become a critical factor, since when throughput of prediction is limiting, the accuracy and speed of the prediction form a joint frontier in the number of accurate predictions that can be generated. Very high confidence predictions in the metagenomic atlas are expected to often be reliable at a resolution sufficient for insight similar to experimentally determined structures, such as into the biochemistry of active sites (61); and for many more proteins where the topology is predicted reliably insight can be obtained into function via remote structural relationships that could not be otherwise detected with sequence.

The emergence of atomic level structure in language models reveals a high resolution picture of protein structure that is encoded by evolution into sequence patterns across millions of proteins, adding to the evidence that the unsupervised training objective materializes deep information about the biology of proteins. ESM-2 is the result of our work over several years focusing on emergence of biological properties, and is the first time a language model has been shown to capture a high resolution picture of structure. Our current models are very far from the limit of scale in parameters, sequence data, and compute that can in principle be applied. We are optimistic that as we continue to scale there will be further emergence. Our results showing the improvement in the modeling of low depth proteins point in this direction.

ESM-2 results in an advance in speed that in practical terms is up to one to two orders of magnitude, which puts far larger numbers of sequences within reach of accurate atomic level structure prediction. Obtaining hundreds of millions of predicted structures within practical timescales can help to reveal new insights into the breadth and diversity of natural proteins, and to accelerate discovery of new protein structures and functions.

## 7. Acknowledgements

We would like to thank FAIR team members Naman Goyal, Yann LeCun, Adam Lerer, Jason Liu, Laurens van der Maaten, Sainbayar Sukhbaatar, and collaborators Justas Dauparas, and Sergey Ovchinnikov, for technical help, feedback, and discussions that helped shape this project. We thank Eugene Koonin and Feng Zhang for feedback on the metagenomic dataset.

We thank Ammar Rizvi, Jon Shepard, and Joe Spisak for program support. We thank Sahir Gomez, Somya Jain, William Ngan, and Neil Seejoor for their work on the ESM Metagenomic Atlas website.

We also thank the developers of the OpenFold project, fairseq, PyTorch, Foldseek, MMseqs2, PyMol, Biotite, and others for building invaluable open source tools, and the creators and maintainers of MGnify, PDB, UniProt, and UniRef, as well as the researchers whose experimental efforts are included in these resources.

There were no external sources of funding for the project. The authors declare no competing financial interests. No patent applications have been filed on this work. Models and data have been released under permissive licenses (MIT license for models and CC-BY for data) in accordance with the principles of open science.

## A. Materials and Methods

### A.1. Data

#### A.1.1. Sequence dataset used to train ESM-2

UniRef50, September 2021 version, is used for the training of ESM models. The training dataset was partitioned by randomly selecting 0.5% (≈ 250,000) sequences to form the validation set. The training set has sequences removed via the procedure described in Meier et al. (54). MMseqs search (`-min-seq-id 0.5 -alignment-mode 3 -max-seqs 300 -s 7 -c 0.8 -cov-mode 0`) is run using the train set as query database and the validation set as target database. All train sequences which match a validation sequence with 50% sequence identity under this search are removed from the train set.

De-novo designed proteins are filtered out from the pretraining dataset via two filters. First, any sequence in UniRef50 and UniRef90 that was annotated as “artificial sequence” by a taxonomy search on the UniProt website, when `2021 04` was the most recent release (1,027 proteins), was removed. Second, jackhmmer was used to remove all hits around a manually curated set of 81 de-novo proteins. jackhmmer was run with `--num-iter 1 -max` flags, with each of the 81 de-novo proteins as a query and UniRef100 as a search database. All proteins returned by jackhmmer were removed from both UniRef50 and UniRef90 via their UniRef IDs (58,462 proteins). This filtering is performed to enable future work evaluating the generalization of language models to de-novo sequences.

To increase the amount of data and its diversity, a minibatch of UniRef50 sequences is sampled for each training update. Each sequence is then replaced with a sequence sampled uniformly from the corresponding UniRef90 cluster. This allowed ESM-2 models to train on over 60M protein sequences.

#### A.1.2. Structure training sets for ESMFold

For training ESMFold, we follow the training procedure outlined in Jumper et al. (12). We find all PDB chains until 2020-05-01 with resolution less than or equal to 9Å and length greater than 20. All proteins where over 20% of the sequence is the same residue is not considered. MM-seqs easy-cluster with default parameters is used to cluster resulting sequences at 40% sequence identity. Only individual chains are used during training, even when the chain is part of a protein complex. This results in 25,450 clusters covering a total of 325,498 chains.

At training time, each cluster is sampled evenly, and then a random protein is sampled from each cluster. Rejection sampling is applied to train on longer proteins more frequently, where protein chains are accepted with probability 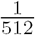 max(min(Nres, 512), 256).

As described in Hsu et al. (62), we generated a set of 13,477,259 structure predictions with AlphaFold2 using MSAs generated via the process in Rao et al. (40). The dataset is then filtered to select only sequences with mean pLDDT *>* 70. Because of the way the dataset is constructed, only 1.5% of the dataset is removed with this filter. Additionally, loss is not calculated for residues with pLDDT *<* 70. We found that this is necessary to obtain increased performance using predicted structures. Predicted structures are sampled 75% of the time, and real structures 25% of the time during training. Data processing is done with Biotite (63).

#### A.1.3. Structure validation and test sets

During method development (e.g. hyperparameter selection), we used a temporally held out validation set obtained from the Continuous Automated Model Evaluation (CAMEO) server (36) by filtering from August 2021 to January 2022.

We report results by testing 3D structure prediction models on two test sets, both chosen to be temporally held out from our supervised training set. The first test set is from CAMEO, consisting of all 194 test proteins from April 01, 2022 through June 25, 2022. Our second test set consists of 51 targets from the CASP14 competition (37). For both test sets, metrics are computed on all modeled residues in the PDB file. The full CASP14 target list is:

T1024, T1025, T1026, T1027, T1028, T1029, T1030, T1031, T1032, T1033, T1034, T1035, T1036s1, T1037, T1038, T1039, T1040, T1041, T1042, T1043, T1044, T1045s1, T1045s2, T1046s1, T1046s2, T1047s1, T1047s2, T1049, T1050, T1053, T1054, T1055, T1056, T1057, T1058, T1064, T1065s1, T1065s2, T1067, T1070, T1073, T1074, T1076, T1078, T1079, T1080, T1082, T1089, T1090, T1091, T1099.

These is the full extent of the publicly available CASP14 targets as of July 2022.

No filtering is performed on these test sets, as ESMFold is able to make predictions on all sequences, including the length-2166 target T1044.

#### A.1.4. CAMEO Dataset Difficulty Categories

The CAMEO evaluation places each target into three categories: easy, medium, and hard. This placement is done based on the average performance of all public structure prediction servers. Targets are classified as “easy” if the average LDDT is *>* 0.75, “hard” if the average LDDT is *<* 0.5, and “medium” otherwise. In the main text, we report average performance across all targets in CAMEO. In Table S4 we provide statistics for each difficulty category.

### A.2. Language Models

#### A.2.1. Computing unsupervised contact prediction from language models

We use the methodology of Rao et al. (34) to measure unsupervised learning of tertiary structure in the form of contact maps. A logistic regression is used to identify contacts. The probability of a contact is defined as

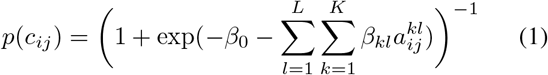

where *c*_*ij*_ is a boolean random variable which is true if amino acids *i, j* are in contact. Suppose our transformer has *L* layers and *K* attention heads per layer. Then *A*_*kl*_ is the symmetrized and APC-corrected (64) attention map for the *k*-th attention head in the *l*-th layer of the transformer, and 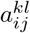 is the value of that attention map at position *i, j*.

The parameters are fit in scikit-learn (65) using L1-regularized logistic regression with *λ* = 0.15. The regression is fit using the same 20 protein training set used in Rao et al. (34), which was simply a random selection from the trRosetta (11) training set. We performed a variability analysis using 20 bootstrapped samples of 20 training proteins from the total set of 14862 proteins. The average long range P@L was 0.4287 with a standard deviation of 0.0028. We also performed experiments using larger training sets, but observed no significant performance change. Given these results, we are confident that selecting a subset of 20 proteins for training provides a good estimate of contact precision performance.

Unsupervised contact prediction results are reported for the 14842 protein test set used in Rao et al. (34), which is also derived from the trRosetta training set, excluding the 20 proteins used in fitting the regression. For both training and test a contact is defined as two amino acids with C-*α* distance *<* 8Å.

#### A.2.2. Language model perplexity calculations

Perplexity is a measure of a language model’s fidelity and is defined as the exponential of the negative log-likelihood of the sequence. Unfortunately, there is no efficient method of computing the log-likelihood of a sequence under a masked language model. Instead, there are two methods we can use for estimating perplexity.

First, let the mask *M* be a random variable denoting a set of tokens from input sequence *x*. Each token has a 15% probability of inclusion. If included the tokens have an 80% probability of being replaced with a mask token, a 10% probability of being replaced with a random token, and a 10% probability of being replaced with an unmasked token. Let 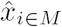 denote the set of modified input tokens. The perplexity is then defined as

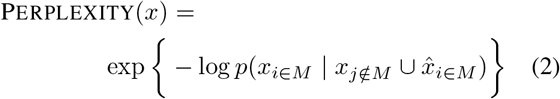

As the set *M* is a random variable, this expression is non-deterministic. This makes it a poor estimate of the perplexity of a single sequence. However, it requires only a single forward pass of the model to compute, so it is possible to efficiently obtain an estimate of the expectation of this expression over a large dataset. When reporting the perplexity over a large dataset (such as our UniRef validation set), this estimate is used.

The second perplexity calculation is the pseudo-perplexity, which is the exponential of the negative pseudo-log-likelihood of a sequence. This estimate provides a deterministic value for each sequence, but requires L forward passes to compute, where L is the length of the input sequence. It is defined as

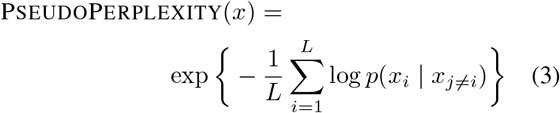

When reporting the perplexity for an individual sequence (e.g. on CASP14 or CAMEO), this estimate is used. For brevity, we refer to both of these estimates as the “perplexity,” as they can be interpreted in a similar manner.

#### A.2.3. ESM-2 model architecture

We use a BERT (18) style encoder only transformer architecture (16) with modifications. We change the number of layers, number of attention heads, hidden size and feed forward hidden size as we scale the ESM model (Table S3).

The original transformer paper uses absolute sinusoidal positional encoding to inform the model about token positions. These positional encodings are added to the input embeddings at the bottom of the encoder stack. In ESM-1b (14), we replaced this static sinusoidal encoding with a learned one. Both static and learned absolute encodings provide the model a very cheap way of adding positional information. However, absolute positional encoding methods don’t extrapolate well beyond the context window they are trained on. In ESM-2, we used Rotary Position Embedding (RoPE) (66) to allow the model extrapolate beyond the context window it is trained on. RoPE slightly increases the computational cost of the model, since it multiplies every query and key vector inside the self attention with a sinusoidal embedding. In our experiments, we observed that this improves model quality for small models. However, we observed that the performance improvements start to disappear as the model size and training duration get bigger.

#### A2.4. Training ESM-2

In ESM-2, we have made multiple small modifications to ESM-1b with the goal of increasing the effective capacity. ESM-1b had dropout both in hidden layers and attention which we removed completely to free up more capacity. In our experiments, we did not observe any significant performance regressions with this change.

We trained most of our models on a network with multiple nodes connected via a network interface. As the models get bigger, the amount of communication becomes the fundamental bottleneck for the training speed. Since BERT style models have been shown to be amenable to very large batch sizes (67), we increased our effective batch size to 2M tokens.

For model training optimization, we used Adam with *β*_1_ = 0.9, *β*_2_ = 0.98, *ϵ* = 10^*−*8^ and *L*_2_ weight decay of 0.01 for all models except the 15 billion parameter model, where we used a weight decay of 0.1. The learning rate is warmed up over the first 2,000 steps to a peak value of 4e-4 (1.6e-4 for the 15B parameter model), and then linearly decayed to one tenth of its peak value over the 90% of training duration. We trained all models for 500K updates except the 15B model which we trained for 270K steps. All models used 2 million tokens as batch size except the 15B model where we used 3.2 million tokens batch size. In order to efficiently process large proteins, we cropped long proteins to random 1024 tokens. We used BOS and EOS tokens to signal the beginning and end of a real protein, to allow the model to separate a full sized protein from a cropped one.

We used standard distributed data parallelism for models up to 650M parameters and used sharded data parallelism (FSDP) (68) for the 2.8B and 15B parameter models. FSDP shards model weights and optimization parameters across multiple GPUs, allowing us to train models that can’t fit into a single GPU memory.

#### A.2.5. ESM-2 ablation experiments

We ran ablation experiments using 150M parameter models trained for 100K steps. Ablations were performed for RoPE, the training dataset (comparing to the ESM-1b training dataset), and UniRef90 sampling (Table S5).

Unsupervised contact prediction results show that both RoPE and newer data significantly improve the results. We do observe a slight regression when sampling from UniRef90 clusters, however we believe this difference is small and the UniRef90 cluster sampling is likely to help for the larger models.

#### A.2.6. Relationship between change in Perplexity and Contact Accuracy

The relationship between improvements in perplexity and improvements in contact accuracy can be measured via normalized discounted cumulative gain (NDCG). In particular, we hypothesize that large improvements in perplexity correspond with large improvements in contact accuracy. We define the change in perplexity as the difference in language model perplexity for a particular protein sequence between adjacent model sizes. Similarly, we define the change in contact accuracy as the difference in unsupervised contact precision for a particular protein sequence between adjacent model sizes. By ranking proteins according to the change in perplexity, we then compute the NDCG with respect to the change in contact accuracy. The average NDCG across the five model classes is 0.87.

### A.3. ESMFold

#### A.3.1. ESMFold model architecture

The ESMFold model uses a simple architecture that leverages the evolutionary information captured by the language model. The architecture is split into two parts, similarly to AlphaFold2: a folding module which takes the language model features as input and produces representations, and a structure module which takes the output from the folding module and outputs 3d atomic coordinates. For the structure module, we use the equivariant transformer architecture with invariant point attention proposed in AlphaFold2. For the folding block we simplify the Evoformer block used in AlphaFold2. No templates are used in ESMFold.

The major change that needs to be made to adapt the Evo-former block to language model features is to remove its dependence on MSAs. Since MSAs are two dimensional, the Evoformer employs axial attention (69) over the columns and rows of the MSA. The language model features are one dimensional, so we can replace the axial attention with a standard attention over this feature space. The self-attention uses a bias derived from the pairwise representations. The sequence representation communicates with pairwise representation via both an outer product and outer difference. Other operations in the Evoformer block are kept the same. We call this simplified architecture the Folding block, described in detail in Algorithm 1, and shown in Fig. 2A.

Our final architecture, ESMFold, described in Algorithm 2, has 48 folding blocks. It is trained for an initial 125K steps on protein crops of size 256, and then fine-tuned with the structural violation loss for 25K steps, on crop sizes of 384. We use the Frame Aligned Point Error (FAPE) and distogram losses introduced in AlphaFold2, as well as heads for predicting LDDT and the pTM score. We omit the masked language modeling loss. Language model parameters are frozen for training ESMFold. We use the 3B parameter ESM-2 language model, the largest model that permits inference on a single GPU.

##### Algorithm 1 Folding block

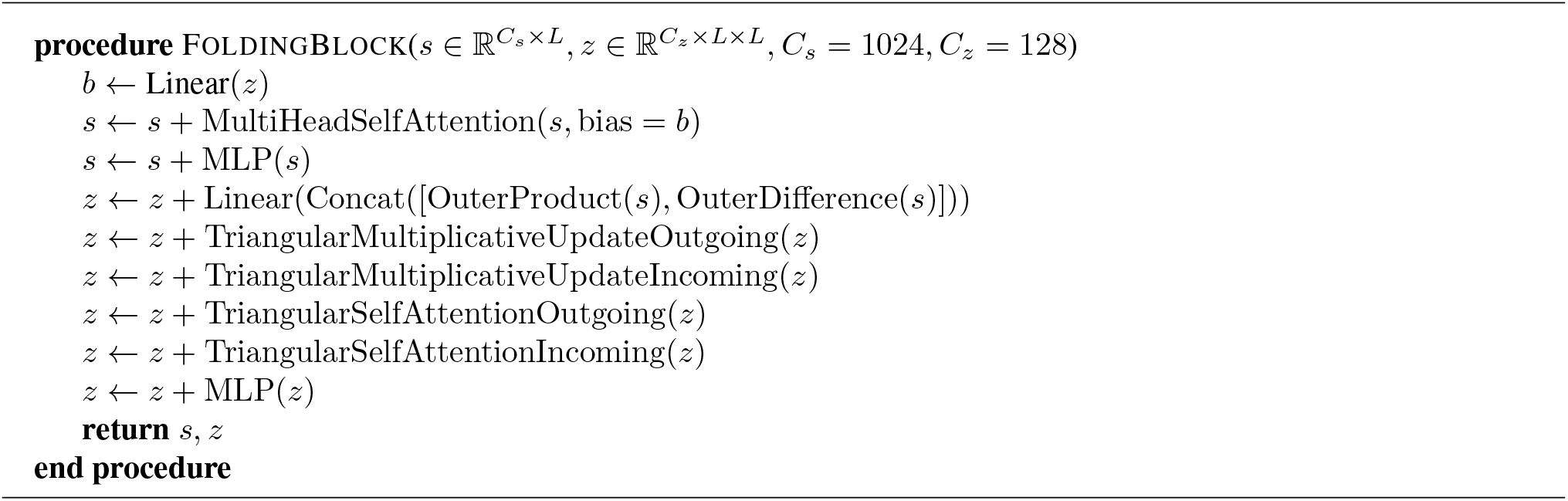

##### Algorithm 2 ESMFold with *N* folding blocks. ESM hiddens returns all hidden representations from an ESM language model. layer weights contains a trainable weight for each layer of ESM

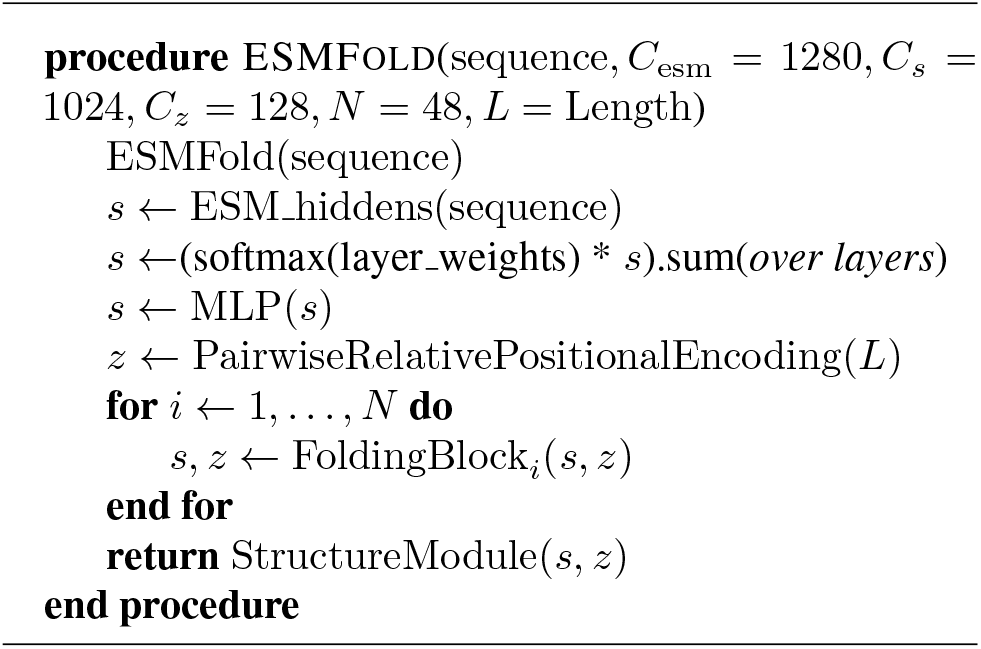

We use a learned weighted sum of ESM embeddings to produce the initial hidden state into the model. This is then fed through an MLP. The initial pairwise state is simply the pairwise relative positional encoding described in Jumper et al. (12). We found that using the attention maps initially gives a boost in performance, but this disappears during training. For experiments that do not use any folding blocks, we use an MLP applied to the ESM attention maps as input, and add the pairwise relative positional encoding to the attention map scores. Finally, the StructureModule projects these representations into coordinates.

The predicted LDDT head is output from the hidden representation of the StructureModule. The predicted TM head uses the pairwise representation z. Finally, we also predict the distogram, from the same representation.

#### A.3.2. Masked prediction

It is possible to sample alternate predictions from ESMFold by masking inputs to the language model. We test this procedure with the following protocol: Input 1000 different sequences into ESMFold with different masking patterns in the language model. The masking patterns are uniformly sampled, where 0 to 15% of the sequence is masked out. A prediction is made for each masked sequence, and the sequence with highest pLDDT is chosen as the final model prediction. On average, applying this procedure only results in a 0.021 LDDT increase on CAMEO, but on some PDBs can substantially improve the accuracy, e.g. for PDB 6s44, TM-score improves from 0.81 to 0.94 (Fig. S6).

#### A.3.3. Extracting coordinates from ESM-2

The following methodology is used to project out coordinates from the language model representations (Fig. 1, Table S1). We train an equivariant structure module directly on top of the frozen ESM representations using a dataset of experimentally determined structures. The training set is the same as used for ESMFold, and we use the same losses and architecture as the AlphaFold2 structure module. We initialize the pairwise representation of the structure module with the output of an MLP that processes the attention maps of the language model. Note that we do not use the predicted structures dataset as data augmentation in these experiments; we train the projection only with experimentally determined structures.

As language models grow in size, we find a large increase in LDDT, from 0.48 on the 8M parameter LM to 0.72 on the 15B parameter LM. This demonstrates that a simple head on top of a powerful language model already gives reasonably accurate structure predictions.

#### A.3.4. Timing analysis

We evaluate the speed of the model by testing sequences of varying length on a single NVIDIA V100 GPU. ESMFold makes a prediction on a protein with 384 residues in 14.2 seconds, 6x faster than a single AlphaFold2 model. On shorter sequences we see a 60x improvement (Fig. S2). Note that this excludes the CPU time for MSA and template search, as well as the 5x from the default ensemble of models used by AlphaFold2. ESMFold can be run reasonably quickly on CPU, and an Apple M1 Macbook Pro makes the same prediction in just over 5 minutes.

ESMFold provides multiple options for reducing GPU memory utilization including chunked attention, mixed precision, and CPU offloading, some of which come at the cost of inference speed. Combined, the optimizations allow predictions on long sequences (such as length-2166 CASP14 target T1044) on an NVIDIA V100 GPU.

### A.4. Metagenomic predictions

#### A.4.1. Folding 617 million sequences from Mgnify

We obtained MGnify (25) version 2022 at 90% sequence similarity (MGnify90). We built a fault tolerant distributed system with a main node which, via TCP, communicates sequences to many workers and receives results as folded protein structures. We were able to leverage the resources of a heterogeneous GPU cluster consisting of P100s, V100s, and A100s of various configurations. We estimate that on a homogeneous network GPU cluster of V100s, the entire 620 million sequences would take approximately 28,000 GPU days to fold, which we were able to do in 2 weeks time. We obtained structure predictions and corresponding pLDDT values for each of these sequences.

#### A.4.2. Analysis of folded metagenomics structures

On a random sample of 1M high confidence structures, we used Foldseek search (version 3.915ef7d) (46) to perform an all-by-all structural similarity search against the PDB (as of April 12, 2022) based on TM-score. We use foldseek with default parameters, except increasing the E-value to 1.0 from the default 1e-3 (foldseek search -e 1.0), to increase recall. We also used MMseqs2 search (version 13.45111) to perform an all-by-all sequence similarity search against UniRef90. We use MMseqs2 with default parameters, except that we re-ran MMseqs2 with the most sensitive setting (-s 7.0) for any sequences that returned an empty result, to increase the recall.

To visualize this landscape of 1M MGnify sequences, we first used ESM-1b to embed each sequence as a 1280-dimensional vector. These embeddings were then visualized using the umap version 0.5.3, scanpy version 1.9.1, and anndata 0.8.0 Python packages (70–72), where dimensionality reduction was applied directly to the embedding vectors (use rep=‘X’ in scanpy.tl.umap) with default parameters (15-nearest-neighbors graph via approximate Euclidean distance, UMAP min dist=0.5).

We further analyzed a random subsample of very high-confidence structures with mean pLDDT greater than 0.9, corresponding to ∼59K structures. For each of these structures, we used Foldseek easy-search (`--alignment-type 1`) to identify similar structures in the PDB. To assess the quality of structure predictions with no Foldseek matches, we used full AlphaFold2 with MSAs to also obtain structure predictions, where we picked the top of five relaxed models ranked by mean pLDDT. We then computed RMSD values of aligned backbone coordinates and all-atom TM-score between the ESMFold-predicted and AlphaFold2-predicted structures and found good agreement of the predictions between both methods (Fig. S7).

To select our case studies, we then used blastp version 2.10.0+ to search for similar sequences in UniRef90 to compute sequence identity. For case-study sequences with no significant matches in UniRef90, we also used the jackhmmer web server (https://www.ebi.ac.uk/Tools/hmmer/search/jackhmmer) (47) to manually query four reference proteomes for similar sequences. Highlighted structure predictions with low similarity to known structures were manually selected and are summarized in Fig. 4. For these structures, we also performed an additional structural similarity search using the Foldseek webserver (https://search.foldseek.com/search) with default parameters to identify the closest structures in PDB100 211201 beyond the TM-score cutoff of 0.5.

### A.5. Multimer Benchmark

#### A.5.1. Recent-PDB-Multimers

To evaluate ESMFold on protein complexes. We construct an evaluation set using the methods described in Evans et al. (45). This dataset consists of targets deposited in the Protein Data Bank between 2020-05-01 and 2022-06-01. The following filtering steps are performed:

- Complexes must contain more than 1 chain and less than 9 chains.
- Chains with length *<* 20 residues, or where one residue makes up *>* 20% of the chain are excluded.
- Complexes must contain fewer than 1536 residues, excluding chains which fail the previous step.
- Each chain is assigned to a 40% overlap cluster using clusters provided by the PDB
- Each complex is assigned a cluster which is the union of chain cluster ids
- From each cluster complex, the example with highest resolution is selected as the representative

These steps result in a total of 2978 clusters. Predictions are made on the full complex, but metrics are computed on a per chain-pair basis using the DockQ program (39). Chain pairs are greedily selected for evaluation if their pair cluster id has not been previously evaluated. Chain pairs which DockQ identifies as having no contacting residues in the ground truth are not evaluated. This results in a total of 3505 unique chain pairs.

#### A.5.2. Multimer Predictions

To predict complexes in the benchmark shown in Figs. 2D and S4, we give a residue index break of 1000 to ESMFold and link chains with a 25-residue poly-glycine linker, which we remove before displaying. Note that this is using ESM-Fold out of distribution since single chains are used during training.

### A.6. Orphan Proteins

Orphan proteins are sequences with few to no evolutionary homologs in either structure or sequence databases. Due to a lack of evolutionary information, these sequences can be very challenging for current structure prediction models. To evaluate ESMFold on orphan proteins, we construct an orphan protein dataset using the following procedure:

- Select structures deposited in the PDB from 2020-05-01 to 2022-05-01 with resolution greater than 9Å and at least 20 modeled residues.
- Cluster at a 70% sequence identity threshold with mmseqs, and select the cluster representatives.
- Run hhblits for 1 iteration (all other parameters default) against UniRef (2020 06), select sequences with no hits.
- Run the standard AlphaFold2 MSA generation pipeline against UniRef, MGnify, and BFD, selecting sequences with *<* 100 total sequence hits and no template hits with TM-score *>* 0.5.

Fig. S8 shows results at different MSA depth thresholds. After filtering, there are 104 sequences with MSA depth ≤ 100, 70 sequences with MSA depth ≤ 10, and 22 sequences with MSA depth = 1. Beyond the constraint that no template has TM-score *>* 0.5, no filtering on the number of templates is performed.

**Figure S1.**
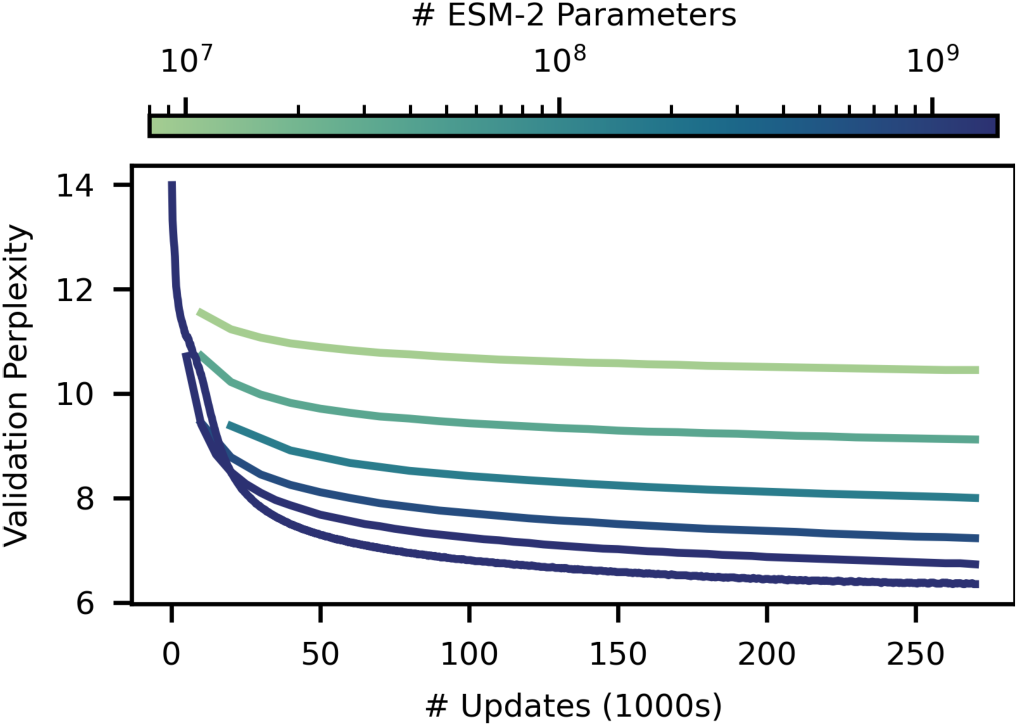
ESM-2 masked language modeling training curves. Training curves for ESM-2 models from 8M (highest curve, light) to 15B parameters (lowest curve, dark). Models are trained to 270K updates. Validation perplexity is measured on a 0.5% random-split holdout of UniRef50. After 270K updates the 8M parameter model has a perplexity of 10.45, and the 15B model reaches a perplexity of 6.37.

**Figure S2.**
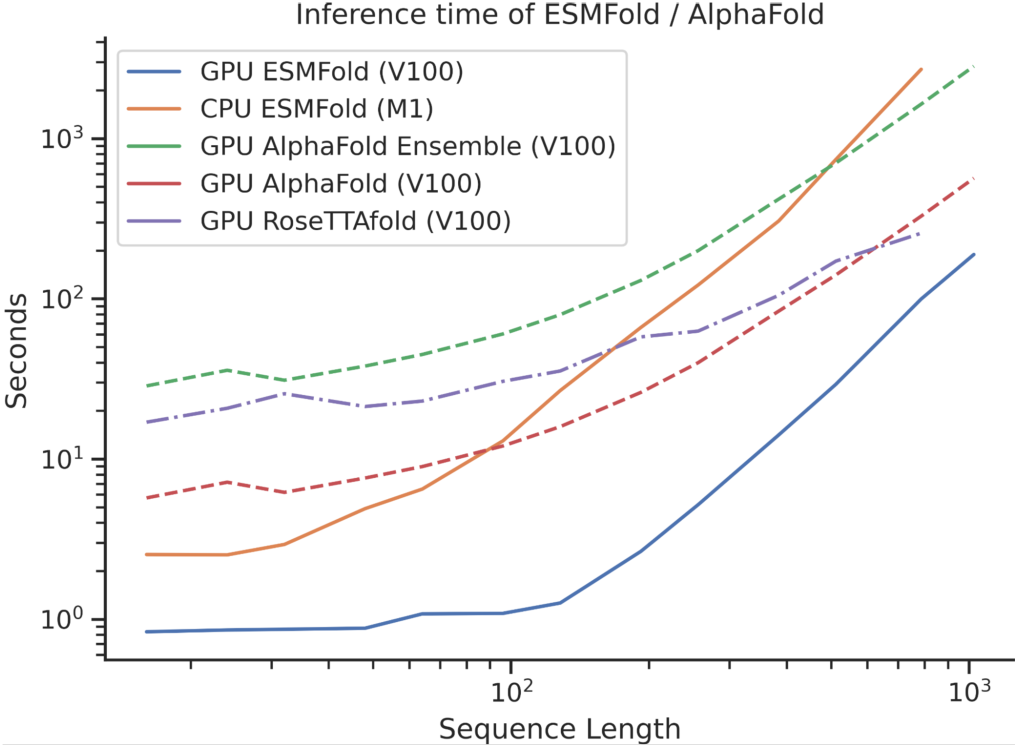
ESMFold timing. Comparison to AlphaFold2 and RoseTTAfold. We test the speed of ESMFold on sequence lengths up to 1024. Note that this comparison is only on the network forward time, and does not include the cost of the search to generate MSAs. ESMFold performance at low sequence lengths is dominated by the forward pass of the language model. At high sequence lengths the *O*(*N* ^3^) computation of pairwise representations takes over. Most of ESMFold’s speed advantage comes from not needing to process the MSA branch. We see an over 60x speed advantage for shorter protein sequences, and a reasonable speed advantage for longer protein sequences. We do not count Jax graph compilation times or MSA search times for AlphaFold2, meaning in practice there is a larger performance difference in the cold start case. We also use an optimized Colabfold 1.3.0 (23) to do speed comparison. No significant optimization has been performed on ESMFold, and we suspect that further gains can be made by optimizing ESMFold as well. For RoseTTAfold, the speed of the SE(3) Transformer dominates, especially at low sequence lengths. The number of SE(3) max-iterations are artificially limited to 20 (default 200) and no MSAs are used as input for these measurements. For RoseTTAfold predictions we do not include the cost of computing sidechains with PyRosetta.

**Figure S3.**
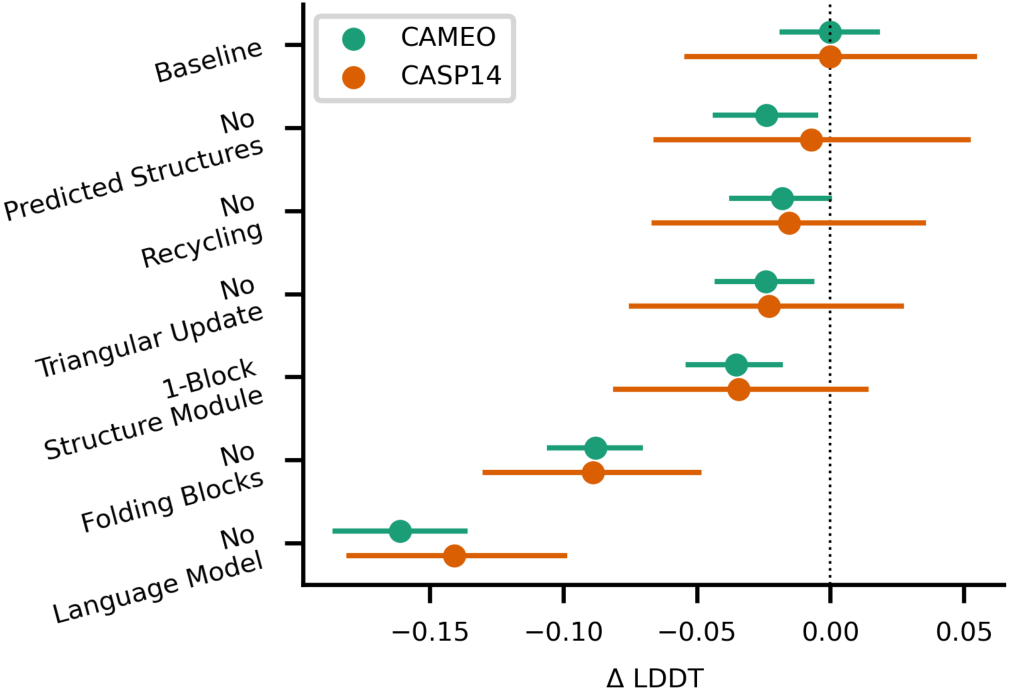
ESMFold ablations on CAMEO and CASP14. ESMFold ablations on CAMEO and CASP14 test sets show the largest contributing factors to performance are the language model and the use of folding blocks. Other ablations reduce performance on CASP14 and CAMEO by 0.01-0.04 LDDT.

**Figure S4.**
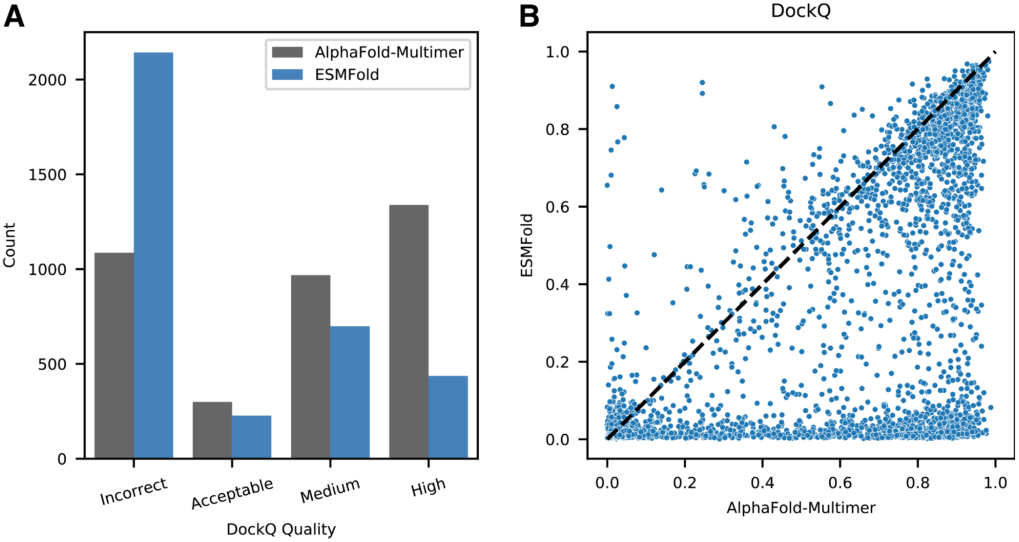
Comparison of ESMFold and AlphaFold-Multimer on recent-PDB-multimers dataset. DockQ (39) scores for AlphaFold-Multimer and ESMFold predictions for chain pairs in the Recent-PDB-Multimers dataset. DockQ qualitative categorizations (left) and quantitative comparison (right) are provided for all chain pairs. ColabFold (23) was used to generate paired MSAs for each complex using the ‘paired+unpaired’ MSA generation setting. UniRef, environmental, and template databases were used. ESMFold predictions are in the same qualitative DockQ categorization for 53.2% of complexes, even though ESMFold is not trained on protein complexes. Dataset generation and scoring methodology described in Appendix A.5.1.

**Figure S5.**
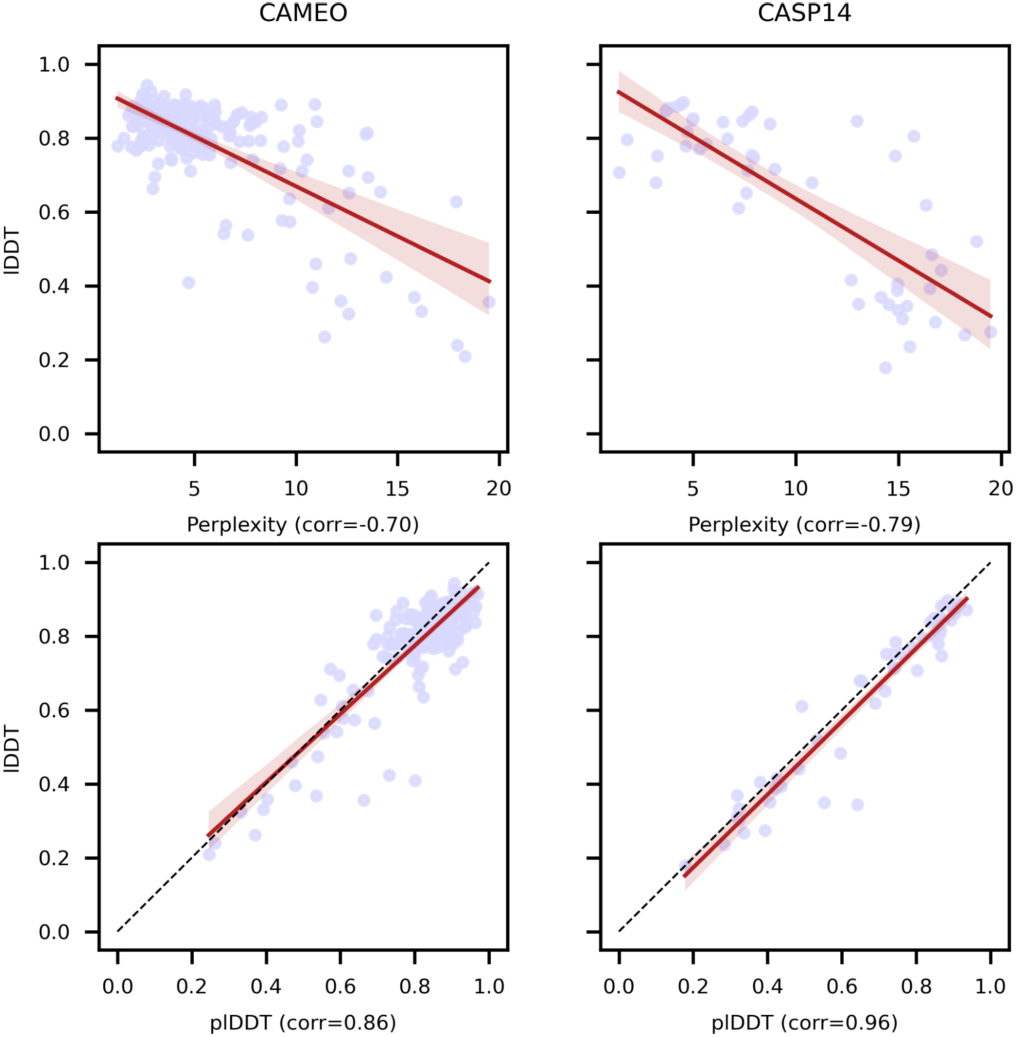
ESMFold calibration with respect to perplexity and pLDDT on CASP14 and CAMEO. Language model perplexity and ESMFold pLDDT are both well correlated with actual structure prediction accuracy on CASP14 and CAMEO. Well understood sequences with language model perplexity *<* 6 are usually well predicted by ESMFold. The strong correlation between pLDDT and LDDT suggests filtering predictions by pLDDT will mostly capture well predicted structures.

**Figure S6.**
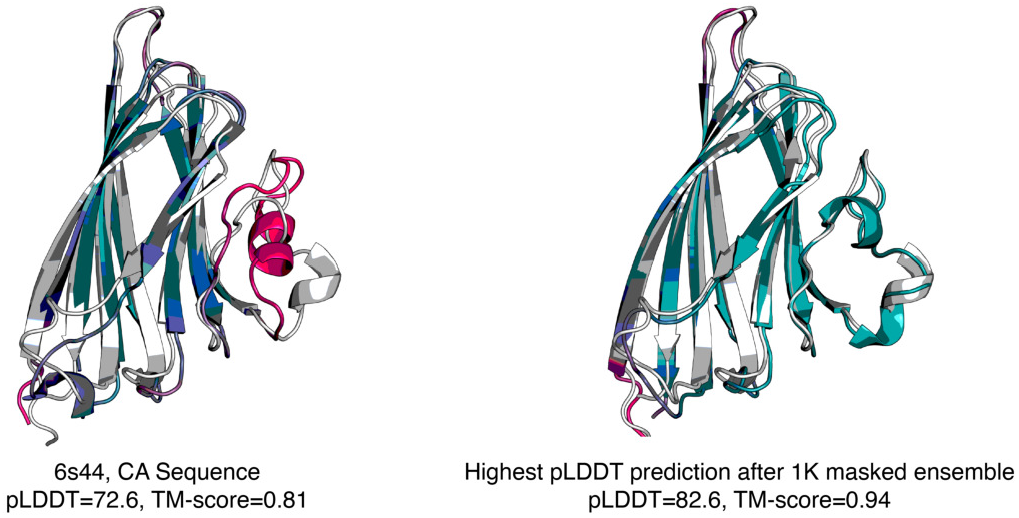
Masked prediction on Cα sequence of PDB 6s44. Left: ESMFold prediction (TM-score=0.81) on the C*α* sequence of PDB 6s44. Right: Best prediction out of 1000 masked sequences generated via the procedure described in Appendix A.3.2. Prediction with highest pLDDT is shown, and has improved TM-score (Tm-score=0.94).

**Figure S7.**
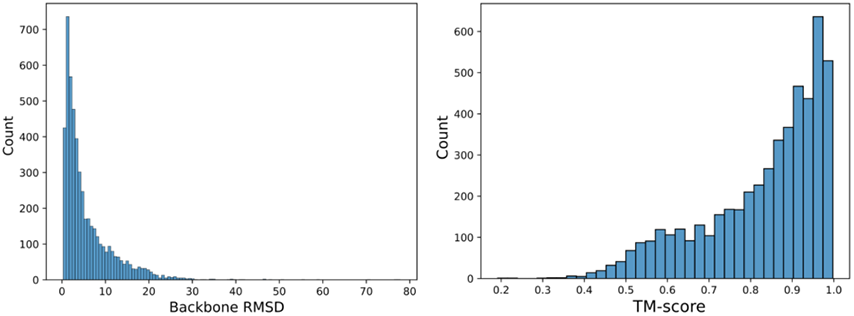
Comparison to AlphaFold2 of structurally remote ESMFold predictions. Distributions of backbone RMSDs (left) and TM-scores (right) of ESMFold-AlphaFold2 predictions of the same sequence, where the ESMFold prediction has both high confidence (mean pLDDT *>* 0.9) and low structural similarity to the PDB (Foldseek closest PDB TM-score *<* 0.5).

**Figure S8.**
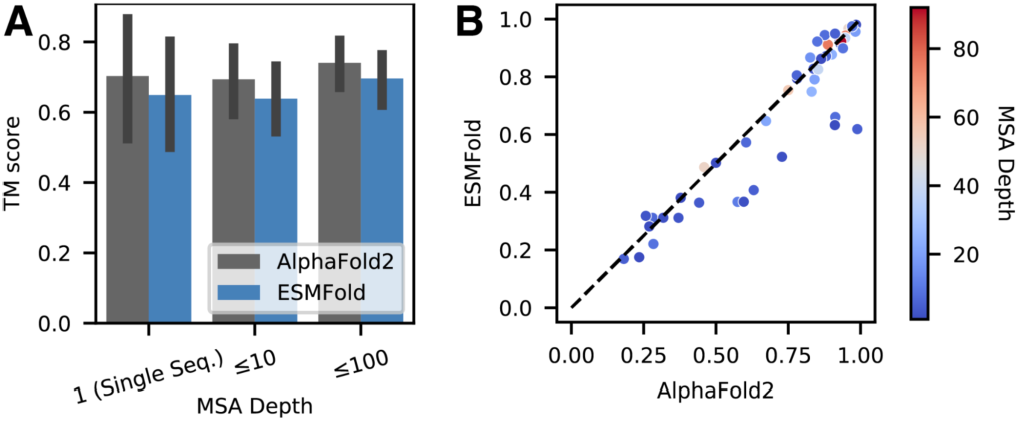
Comparison of ESMFold and AlphaFold2 on a set of orphan proteins. Performance of ESMFold and AlphaFold2 on a set of “orphan proteins” - sequences with few sequence or structural homologs. All compared sequences are temporally held out from the training set. The standard AlphaFold2 sequence and template search pipeline is used to find homologs (dataset construction described in Appendix A.6). (A) Comparison on natural proteins with various MSA depths. Depth is the total number of hits across UniRef and metagenomic databases. (B) TM-score comparison of all individual orphans.

**Table S1.** Detailed language model comparison. Comparison at different numbers of parameters and at different numbers of training updates. Training updates and validation perplexity are not reported for baseline models, since there is no straightforward comparison. For the number of training updates, different models use different batch sizes, so the number of sequences seen can vary even if the number of updates are the same. For validation perplexity, baseline models are not trained on the same dataset, and do not share a common heldout validation set with ESM-2. Prot-T5 is an encoder-decoder language model. Only the encoder portion of the model was used in this evaluation, however the number of parameters reported is the total number of parameters used for training. Unsupervised contact precision results, in the form of long range precision at L and at L / 5, do allow us to compare all transformer language models despite variance in training data. However, CARP, a convolution based language model, does not have attention maps. Note: ESM-1b is evaluated only on sequences of length *<* 1024, due to constraints with position embedding.

| Model | # Params | # Updates | Validation Perplexity | LR P@L | LR P@L/5 | CASP14 | CAMEO |
| --- | --- | --- | --- | --- | --- | --- | --- |
| ESM-2 | 8M | 270K | 10.45 | 0.16 | 0.28 | 0.37 | 0.48 |
|  | 35M | 270K | 9.12 | 0.29 | 0.49 | 0.41 | 0.56 |
|  | 150M | 270K | 8.00 | 0.42 | 0.68 | 0.47 | 0.63 |
|  | 650M | 270K | 7.23 | 0.50 | 0.77 | 0.51 | 0.68 |
|  | 3B | 270K | 6.73 | 0.53 | 0.80 | 0.51 | 0.71 |
|  | 8M | 500K | 10.33 | 0.17 | 0.29 | 0.37 | 0.48 |
|  | 35M | 500K | 8.95 | 0.30 | 0.51 | 0.41 | 0.56 |
|  | 150M | 500K | 7.75 | 0.44 | 0.70 | 0.49 | 0.65 |
|  | 650M | 500K | 6.95 | 0.52 | 0.79 | 0.51 | 0.70 |
|  | 3B | 500K | 6.49 | **0.54** | 0.81 | 0.52 | **0.72** |
|  | 15B | 270K | **6.37** | **0.54** | **0.82** | **0.55** | **0.72** |
| ESM-1b | 650M | — | — | 0.41 | 0.66 | 0.42 | 0.64 |
| Prot-T5-XL (UR50) (21) | 3B | — | — | 0.48 | 0.72 | 0.50 | 0.69 |
| Prot-T5-XL (BFD) (21) | 3B | — | — | 0.36 | 0.58 | 0.46 | 0.63 |
| CARP (24) | 640M | — | — | — | — | 0.42 | 0.59 |

**Table S2.**
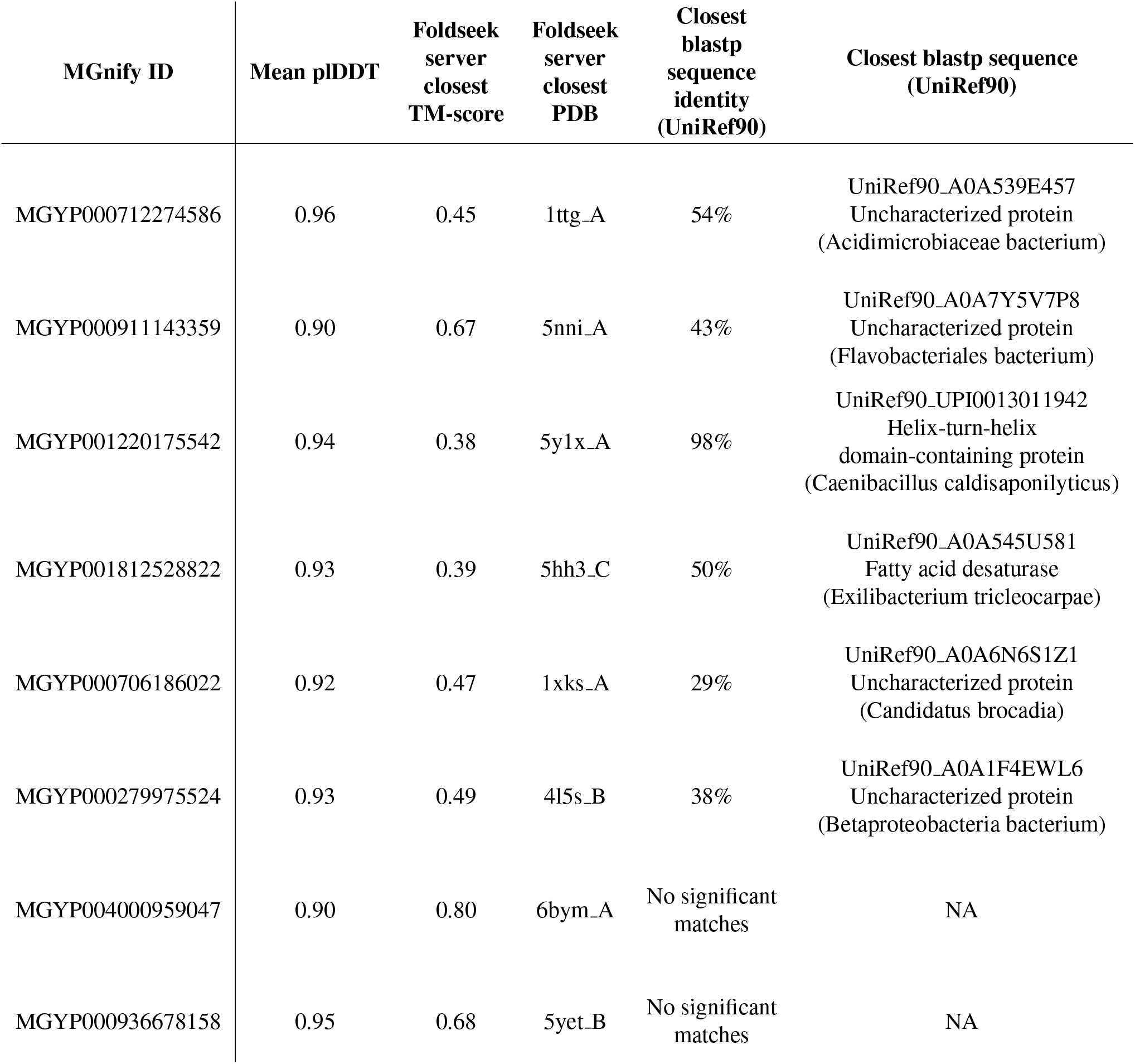
Information on highlighted MGnify proteins. MGnify sequence identifiers corresponding to predicted structures highlighted throughout this study, including the PDB chain and corresponding TM-score of the closest structure identified by the Foldseek webserver as well as the UniRef90 entry and sequence identity of the closest sequence identified by blastp (Appendix A.4.2).

**Table S3.**
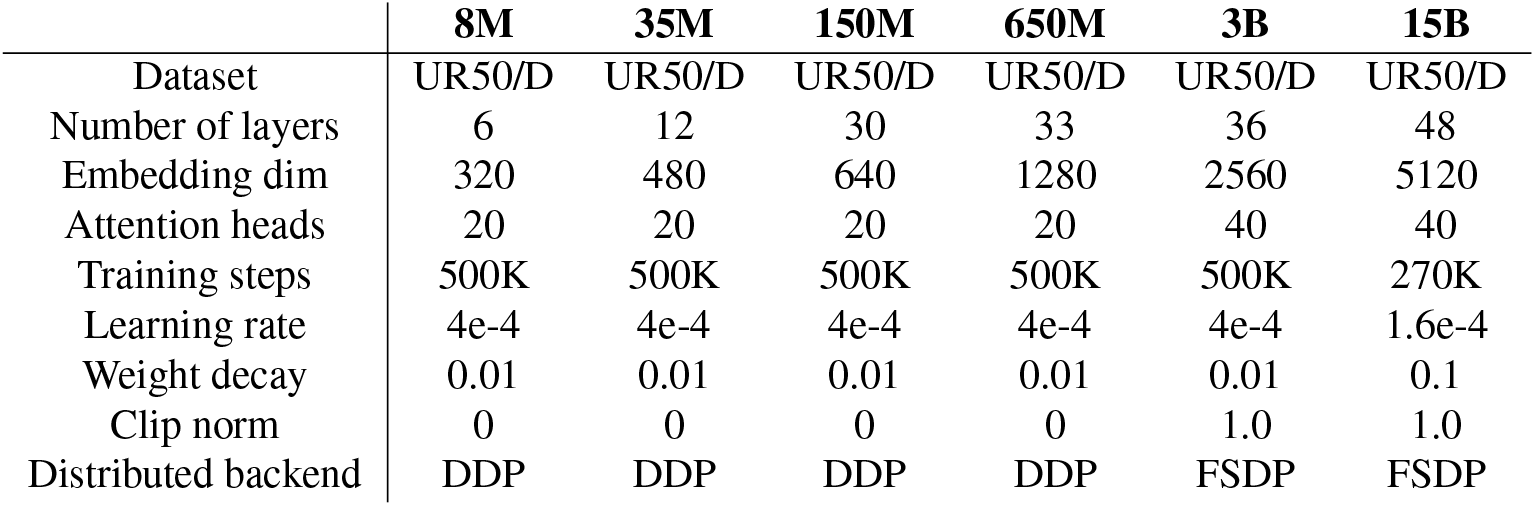
ESM-2 model parameters at different scales.

|  | 8M | 35M | 150M | 650M | 3B | 15B |
| --- | --- | --- | --- | --- | --- | --- |
| Dataset | UR50/D | UR50/D | UR50/D | UR50/D | UR50/D | UR50/D |
| Number of layers | 6 | 12 | 30 | 33 | 36 | 48 |
| Embedding dim | 320 | 480 | 640 | 1280 | 2560 | 5120 |
| Attention heads | 20 | 20 | 20 | 20 | 40 | 40 |
| Training steps | 500K | 500K | 500K | 500K | 500K | 270K |
| Learning rate | 4e-4 | 4e-4 | 4e-4 | 4e-4 | 4e-4 | 1.6e-4 |
| Weight decay | 0.01 | 0.01 | 0.01 | 0.01 | 0.01 | 0.1 |
| Clip norm | 0 | 0 | 0 | 0 | 1.0 | 1.0 |
| Distributed backend | DDP | DDP | DDP | DDP | FSDP | FSDP |

**Table S4.** CAMEO dataset statistics broken down by difficulty class. Median MSA depth is reported for each difficulty class of the CAMEO dataset, along with mean TM-score for ESMFold, AlphaFold, and RoseTTAFold. Half of the samples from the CAMEO dataset consist of “easy” examples, which are well predicted by all models. Differentiation is greater in the “medium” and “hard” classes, which have lower MSA depth and are better predicted by AlphaFold2. Statistics for CASP14 are provided as a comparison. MSA depth numbers provided are from the AlphaFold2 MSA generation pipeline.

| Dataset | Split | Count | MSA Depth<br>(Total) | MSA Depth<br>(UniRef) | ESMFold | AlphaFold2 | RoseTTAFold |
| --- | --- | --- | --- | --- | --- | --- | --- |
| CAMEO | easy | 97 | 21,458 | 17,627 | 0.90 | 0.93 | 0.89 |
|  | medium | 89 | 3,032 | 860 | 0.79 | 0.86 | 0.76 |
|  | hard | 8 | 328.5 | 56 | 0.45 | 0.62 | 0.49 |
| CASP14 | — | 51 | 1228 | 161 | 0.68 | 0.85 | 0.81 |

**Table S5.** ESM-2 architecture ablations.

|  | LR P@L | LR P@L/5 | Validation Perplexity |
| --- | --- | --- | --- |
| Baseline | 0.381 | 0.626 | 8.42 |
| No RoPE | 0.365 | 0.599 | 8.62 |
| Older UniRef Data | 0.368 | 0.599 | 7.98 |
| No UR90 Sampling | 0.387 | 0.631 | 8.40 |

## References

[1] C Yanofsky, V Horn, and D Thorpe. Protein Structure Relationships Revealed By Mutational Analysis. Science (New York, N.Y.), 146(3651):1593–4, December 1964. ISSN 0036-8075. URL http://www.ncbi.nlm.nih.gov/pubmed/14224506.

[2] D Altschuh, T Vernet, P Berti, D Moras, and K Na-gai. Coordinated amino acid changes in homolo-gous protein familiesa. Protein Engineering, Design and Selection, 2(3):193–199, 1988. ISSN 0269-2139. URL http://www.ncbi.nlm.nih.gov/_pubmed/3237684. Publisher: Oxford University Press.

[3] Ulrike Gö bel, Chris Sander, Reinhard Schneider, and Alfonso Valencia. Correlated mutations and residue contacts in proteins. Proteins: Structure, Function, and Bioinformatics, 18(4):309–317, 1994. ISSN 1097-0134. doi: 10.1002/prot.340180402. URL https://onlinelibrary.wiley.com/doi/abs/10.1002/prot.340180402. eprint: https://onlinelibrary.wiley.com/doi/pdf/10.1002/prot.340180402.

[4] Alan S. Lapedes, Bertrand G. Giraud, LonChang Liu, and Gary D. Stormo. Correlated Mutations in Models of Protein Sequences: Phylogenetic and Structural Effects. Lecture Notes-Monograph Se-ries, 33:236–256, 1999. ISSN 07492170. doi: 10.2307/4356049. URL http://www.jstor.org/stable/4356049. Publisher: Institute of Mathe-matical Statistics.

[5] John Thomas, Naren Ramakrishnan, and Chris Bailey-Kellogg. Graphical models of residue coupling in protein families, April 2008. URL https://spubmed.ncbi.nlm.nih.gov/18451428/. ISSN: 15455963 Issue: 2 Pages: 183–197 Publication Title: IEEE/ACM Transactions on Computational Biology and Bioinformatics Volume: 5.

[6] Martin Weigt, Robert A. White, Hendrik Szurmant, James A. Hoch, and Terence Hwa. Identification of direct residue contacts in protein-protein inter-action by message passing. Proceedings of the National Academy of Sciences of the United States of America, 106(1):67–72, January 2009. ISSN 00278424. doi: 10.1073/pnas.0805923106. URL https://www.pnas.org/content/106/1/67https://www.pnas.org/content/106/1/67.abstract. Publisher: National Academy of Sciences.

[7] Faruck Morcos, Andrea Pagnani, Bryan Lunt, Arianna Bertolino, Debora S. Marks, Chris Sander, Riccardo Zecchina, José N. Onuchic, Terence Hwa, and Martin Weigt. Direct-coupling analysis of residue coevolution captures native contacts across many protein families. Proceedings of the National Academy of Sciences of the United States of America, 108(49):E1293–E1301, December 2011. ISSN 00278424. doi: 10.1073/pnas.1111471108. URL https://www.pnas.org/content/108/49/E1293. Publisher: National Academy of Sciences.

[8] Sheng Wang, Siqi Sun, Zhen Li, Renyu Zhang, and Jinbo Xu. Accurate De Novo Prediction of Pro-tein Contact Map by Ultra-Deep Learning Model. PLOS Computational Biology, 13(1):1–34, January 2017. ISSN 1553-7358. doi: 10.1371/journal.pcbi.1005324. URL https://dx.plos.org/10.1371/journal.pcbi.1005324. Publisher: Public Library of Science.

[9] Yang Liu, Perry Palmedo, Qing Ye, Bonnie Berger, and Jian Peng. Enhancing Evolutionary Couplings with Deep Convolutional Neural Net-works. Cell Systems, 6(1):65–74, January 2018. ISSN 24054720. doi: 10.1016/j.cels.2017.11.014. URL https://pubmed.ncbi.nlm.nih.gov/29275173/. Publisher: Cell Press.

[10] John Jumper, Richard Evans, Alexander Pritzel, Tim Green, Michael Figurnov, Kathryn Tunyasuvu-nakool, Olaf Ronneberger, Russ Bates, Augustin Žídek, Alex Bridgland, Clemens Meyer, Simon A A Kohl, Anna Potapenko, Andrew J Ballard, An-drew Cowie, Bernardino Romera-Paredes, Stanislav Nikolov, Rishub Jain, Jonas Adler, Trevor Back, Stig Petersen, David Reiman, Martin Steinegger, Michalina Pacholska, David Silver, Oriol Vinyals, Andrew W Se-nior, Koray Kavukcuoglu, Pushmeet Kohli, and Demis Hassabis. High Accuracy Protein Structure Prediction Using Deep Learning. In Fourteenth Critical Assess-ment of Techniques for Protein Structure Prediction (Abstract Book), page 22. 2020.

[11] Jianyi Yang, Ivan Anishchenko, Hahnbeom Park, Zhenling Peng, Sergey Ovchinnikov, and David Baker. Improved protein structure prediction using predicted interresidue orientations. Proceedings of the National Academy of Sciences, 117(3):1496–1503, Jan-uary 2020. ISSN 0027-8424, 1091-6490. doi: 10.1073/pnas.1914677117. URL https://www.pnas.org/content/117/3/1496. Publisher: National Academy of Sciences Section: Biological Sciences.

[12] John Jumper, Richard Evans, Alexander Pritzel, Tim Green, Michael Figurnov, Olaf Ronneberger, Kathryn Tunyasuvunakool, Russ Bates, Augustin Žídek, Anna Potapenko, Alex Bridgland, Clemens Meyer, Simon A. A. Kohl, Andrew J. Ballard, Andrew Cowie, Bernardino Romera-Paredes, Stanislav Nikolov, Rishub Jain, Jonas Adler, Trevor Back, Stig Petersen, David Reiman, Ellen Clancy, Michal Zielinski, Martin Steinegger, Michalina Pacholska, Tamas Berghammer, Sebastian Bodenstein, David Silver, Oriol Vinyals, Andrew W. Senior, Koray Kavukcuoglu, Pushmeet Kohli, and Demis Hassabis. Highly accurate protein structure prediction with AlphaFold. Nature, 596(7873):583–589, August 2021. ISSN 1476-4687. doi: 10.1038/s41586-021-03819-2. URL https://www.nature.com/articles/s41586-021-03819-2. Bandiera abtest: a Cc license type: cc by Cg type: Nature Research Journals Number: 7873 Primary atype: Research Publisher: Nature Publishing Group Subject term: Com-putational biophysics;Machine learning;Protein structure predictions;Structural biology Subject term id: computational-biophysics;machine-learning;proteinstructure-predictions;structural-biology.

[13] Minkyung Baek, Frank DiMaio, Ivan Anishchenko, Justas Dauparas, Sergey Ovchinnikov, Gyu Rie Lee, Jue Wang, Qian Cong, Lisa N. Kinch, R. Dustin Schaeffer, Claudia Millán, Hahnbeom Park, Carson Adams, Caleb R. Glassman, Andy DeGiovanni, Jose H. Pereira, Andria V. Rodrigues, Alberdina A. van Dijk, Ana C. Ebrecht, Diederik J. Opperman, Theo Sagmeister, Christoph Buhlheller, Tea PavkovKeller, Manoj K. Rathinaswamy, Udit Dalwadi, Calvin K. Yip, John E. Burke, K. Christopher Garcia, Nick V. Grishin, Paul D. Adams, Randy J. Read, and David Baker. Accurate prediction of protein structures and interactions using a three-track neural network. Science, 373(6557):871–876, August 2021. ISSN 0036-8075, 1095-9203. doi: 10.1126/science.abj8754. URL https://www.science.org/doi/10.1126/science.abj8754.

[14] Alexander Rives, Joshua Meier, Tom Sercu, Siddharth Goyal, Zeming Lin, Jason Liu, Demi Guo, Myle Ott, C. Lawrence Zitnick, Jerry Ma, and Rob Fergus. Biological structure and function emerge from scaling unsupervised learning to 250 million protein sequences. Proceedings of the National Academy of Sciences, 118(15), April 2021. ISSN 0027-8424, 1091-6490. doi: 10.1073/pnas.2016239118. URL https://www.pnas.org/content/118/15/e2016239118. Publisher: National Academy of Sciences Section: Biological Sciences.

[15] C. E. Shannon. Prediction and entropy of printed English. The Bell System Technical Journal, 30(1): 50–64, January 1951. ISSN 0005-8580. doi: 10.1002/j.1538-7305.1951.tb01366.x. Conference Name: The Bell System Technical Journal.

[16] Ashish Vaswani, Noam Shazeer, Niki Parmar, Jakob Uszkoreit, Llion Jones, Aidan N Gomez, Łukasz Kaiser, and Illia Polosukhin. Attention Is All You Need. In Advances in Neural Information Processing Systems, pages 5998–6008, 2017. URL https://papers.nips.cc/paper/7181-attention-is-all-you-need.pdf.

[17] Alec Radford, Karthik Narasimhan, Tim Salimans, and Ilya Sutskever. Improving language understanding by generative pre-training. 2018.

[18] Jacob Devlin, Ming-Wei Chang, Kenton Lee, and Kristina Toutanova. BERT: Pre-training of Deep Bidirectional Transformers for Language Understanding. In Proceedings of the 2019 Conference of the North {A}merican Chapter of the Association for Computational Linguistics: Human Language Technologies, Volume 1 (Long and Short Papers), pages 4171–4186, Minneapolis, Minnesota, June 2019. Association for Computational Linguistics. doi: 10.18653/v1/N19-1423. URL http://arxiv.org/abs/1810.04805.

[19] Tom B. Brown, Benjamin Mann, Nick Ryder, Melanie Subbiah, Jared Kaplan, Prafulla Dhariwal, Arvind Neelakantan, Pranav Shyam, Girish Sastry, Amanda Askell, Sandhini Agarwal, Ariel Herbert-Voss, Gretchen Krueger, Tom Henighan, Rewon Child, Aditya Ramesh, Daniel M. Ziegler, Jeffrey Wu, Clemens Winter, Christopher Hesse, Mark Chen, Eric Sigler, Mateusz Litwin, Scott Gray, Benjamin Chess, Jack Clark, Christopher Berner, Sam McCan-dlish, Alec Radford, Ilya Sutskever, and Dario Amodei. Language Models are Few-Shot Learners. CoRR, abs/2005.14165, 2020. URL https://arxiv.org/abs/2005.14165. xeprint: 2005.14165.

[20] Jason Wei, Maarten Bosma, Vincent Y Zhao, Kelvin Guu, Adams Wei Yu, Brian Lester, Nan Du, Andrew M Dai, and Quoc V Le. Finetuned Language Models Are Zero-Shot Learners. page 46, 2022.

[21] Jason Wei, Xuezhi Wang, Dale Schuurmans, Maarten Bosma, Brian Ichter, Fei Xia, Ed Chi, Quoc Le, and Denny Zhou. Chain of Thought Prompting Elicits Reasoning in Large Language Models, une 2022. URL http://arxiv.org/abs/2201.11903. arXiv:2201.11903 [cs].

[22] Aakanksha Chowdhery, Sharan Narang, Jacob De-vlin, Maarten Bosma, Gaurav Mishra, Adam Roberts, Paul Barham, Hyung Won Chung, Charles Sutton, Sebastian Gehrmann, Parker Schuh, Kensen Shi, Sasha Tsvyashchenko, Joshua Maynez, Abhishek Rao, Parker Barnes, Yi Tay, Noam Shazeer, Vinodkumar Prabhakaran, Emily Reif, Nan Du, Ben Hutchin-son, Reiner Pope, James Bradbury, Jacob Austin, Michael Isard, Guy Gur-Ari, Pengcheng Yin, Toju Duke, Anselm Levskaya, Sanjay Ghemawat, Sunipa Dev, Henryk Michalewski, Xavier Garcia, Vedant Misra, Kevin Robinson, Liam Fedus, Denny Zhou, Daphne Ippolito, David Luan, Hyeontaek Lim, Bar-ret Zoph, Alexander Spiridonov, Ryan Sepassi, David Dohan, Shivani Agrawal, Mark Omernick, Andrew M. Dai, Thanumalayan Sankaranarayana Pillai, Marie Pel-lat, Aitor Lewkowycz, Erica Moreira, Rewon Child, Oleksandr Polozov, Katherine Lee, Zongwei Zhou, Xuezhi Wang, Brennan Saeta, Mark Diaz, Orhan Firat, Michele Catasta, Jason Wei, Kathy Meier-Hellstern, Douglas Eck, Jeff Dean, Slav Petrov, and Noah Fiedel. PaLM: Scaling Language Modeling with Pathways, April 2022. URL http://arxiv.org/abs/2204.02311. arXiv:2204.02311 [cs].

[23] Milot Mirdita, Konstantin Schü tze, Yoshitaka Mori-waki, Lim Heo, Sergey Ovchinnikov, and Mar-tin Steinegger. ColabFold: making protein folding accessible to all. Nature Methods, 19 (6):679–682, June 2022. ISSN 1548-7091, 1548-7105. doi: 10.1038/s41592-022-01488-1. URL https://www.nature.com/articles/s41592-022-01488-1.

[24] Martin Steinegger, Milot Mirdita, and Johannes Sö ding. Protein-level assembly increases protein sequence recovery from metagenomic samples manyfold. Nature Methods, 16(7):603–606, 2019. ISSN 1548-7105. doi: 10.1038/s41592-019-0437-4. URL https://doi.org/10.1038/s41592-019-0437-4.

[25] Alex L Mitchell, Alexandre Almeida, Martin Beracochea, Miguel Boland, Josephine Burgin, Guy Cochrane, Michael R Crusoe, Varsha Kale, Simon C Potter, Lorna J Richardson, Ekaterina Sakharova, Maxim Scheremetjew, Anton Korobeynikov, Alex Shlemov, Olga Kunyavskaya, Alla Lapidus, and Robert D Finn. MGnify: the microbiome analysis resource in 2020. Nucleic Acids Research, page gkz1035, November 2019. ISSN 0305-1048, 1362-4962. doi: 10.1093/nar/gkz1035. URL https://academic.oup.com/nar/advance-article/doi/10.1093/nar/gkz1035/5614179.

[26] Supratim Mukherjee, Dimitri Stamatis, Jon Bertsch, Galina Ovchinnikova, Jagadish Chandrabose Sundaramurthi, Janey Lee, Mahathi Kandimalla, I.-Min A. Chen, Nikos C. Kyrpides, and T. B. K. Reddy. Genomes OnLine Database (GOLD) v.8: overview and updates. Nucleic Acids Research, 49(D1):D723–D733, January 2021. ISSN 1362-4962. doi: 10.1093/nar/gkaa983.

[27] Kathryn Tunyasuvunakool, Jonas Adler, Zachary Wu, Tim Green, Michal Zielinski, Augustin Žídek, Alex Bridgland, Andrew Cowie, Clemens Meyer, Agata Laydon, Sameer Velankar, Gerard J. Kleywegt, Alex Bateman, Richard Evans, Alexander Pritzel, Michael Figurnov, Olaf Ronneberger, Russ Bates, Simon A. A. Kohl, Anna Potapenko, Andrew J. Ballard, Bernardino Romera-Paredes, Stanislav Nikolov, Rishub Jain, Ellen Clancy, David Reiman, Stig Petersen, Andrew W. Senior, Koray Kavukcuoglu, Ewan Birney, Pushmeet Kohli, John Jumper, and Demis Hassabis. Highly accurate protein structure prediction for the human proteome. Nature, 596(7873):590–596, August 2021. ISSN 1476-4687. doi: 10.1038/s41586-021-03828-1. URL https://www.nature.com/articles/s41586-021-03828-1. Number: 7873 Publisher: Nature Publishing Group.

[28] Mihaly Varadi, Stephen Anyango, Mandar Deshpande, Sreenath Nair, Cindy Natassia, Galabina Yordanova, David Yuan, Oana Stroe, Gemma Wood, Agata Laydon, Augustin Žídek, Tim Green, Kathryn Tunyasuvunakool, Stig Petersen, John Jumper, Ellen Clancy, Richard Green, Ankur Vora, Mira Lutfi, Michael Figurnov, Andrew Cowie, Nicole Hobbs, Pushmeet Kohli, Gerard Kleywegt, Ewan Birney, Demis Hassabis, and Sameer Velankar. AlphaFold Protein Structure Database: massively expanding the structural coverage of protein-sequence space with high-accuracy models. Nucleic Acids Research, 50(D1):D439–D444, January 2022. ISSN 0305-1048, 1362-4962. doi: 10.1093/nar/gkab1061. URL https://academic.oup.com/nar/article/50/D1/D439/6430488.

[29] Osamu Shimomura, Frank H. Johnson, and Yo Saiga. Extraction, purification and properties of aequorin, a bioluminescent protein from the luminous hydromedusan, aequorea. Journal of Cellular and Comparative Physiology, 59(3):223–239, 1962. doi: https://doi.org/10.1002/jcp.1030590302. URL https://onlinelibrary.wiley.com/doi/abs/10.1002/jcp.1030590302.

[30] K. Mullis, F. Faloona, S. Scharf, R. Saiki, G. Horn, and H. Erlich. Specific enzymatic amplification of DNA in vitro: the polymerase chain reaction. Cold Spring Harbor Symposia on Quantitative Biology, 51 Pt 1:263–273, 1986. ISSN 0091-7451. doi: 10.1101/sqb.1986.051.01.032.

[31] Martin Jinek, Krzysztof Chylinski, Ines Fonfara, Michael Hauer, Jennifer A. Doudna, and Emmanuelle Charpentier. A programmable dual-RNA-guided DNA endonuclease in adaptive bacterial immunity. Science (New York, N.Y.), 337(6096):816–821, August 2012. ISSN 1095-9203. doi: 10.1126/science.1225829.

[32] Baris E Suzek, Yuqi Wang, Hongzhan Huang, Peter B McGarvey, Cathy H Wu, and UniProt Consortium. UniRef clusters: a comprehensive and scalable alternative for improving sequence similarity searches. Bioinformatics, 31(6):926–932, 2014. Publisher: Oxford University Press.

[33] Jesse Vig, Ali Madani, Lav R. Varshney, Caiming Xiong, Richard Socher, and Nazneen Rajani. BERTology Meets Biology: Interpreting Attention in Protein Language Models. September 2020. URL https://openreview.net/forum?id=YWtLZvLmud7.

[34] Roshan Rao, Joshua Meier, Tom Sercu, Sergey Ovchinnikov, and Alexander Rives. Transformer protein language models are unsupervised structure learners. In International Conference on Learning Representations, page 2020.12.15.422761. Cold Spring Harbor Laboratory, December 2021. doi: 10.1101/2020.12.15.422761.

[35] Stephen K Burley, Helen M Berman, Charmi Bhikadiya, Chunxiao Bi, Li Chen, Luigi Di Costanzo, Cole Christie, Ken Dalenberg, Jose M Duarte, Shuchismita Dutta, Zukang Feng, Sutapa Ghosh, David S Goodsell, Rachel K Green, Vladimir Guranoví, Dmytro Guzenko, Brian P Hudson, Tara Kalro, Yuhe Liang, Robert Lowe, Harry Namkoong, Ezra Peisach, Irina Periskova, Andreas Prlí, Chris Randle, Alexander Rose, Peter Rose, Raul Sala, Monica Sekharan, Chenghua Shao, Lihua Tan, Yi-Ping Tao, Yana Valasatava, Maria Voigt, John Westbrook, Jesse Woo, Huanwang Yang, Jasmine Young, Marina Zhuravleva, and Christine Zardecki. RCSB Protein Data Bank: biological macromolecular structures enabling research and education in fundamental biology, biomedicine, biotechnology and energy. Nucleic Acids Research, 47, 2019. doi: 10.1093/nar/gky1004. URL https://academic.oup.com/nar/article-abstract/47/D1/D464/5144139.

[36] Jü rgen Haas, Alessandro Barbato, Dario Behringer, Gabriel Studer, Steven Roth, Martino Bertoni, Khaled Mostaguir, Rafal Gumienny, and Torsten Schwede. Continuous Automated Model EvaluatiOn (CAMEO) complementing the critical assessment of structure prediction in CASP12. Proteins: Structure, Function and Bioinformatics, 86(Suppl 1):387–398, March 2018. ISSN 10970134. doi: 10.1002/prot.25431. Publisher: John Wiley and Sons Inc.

[37] Andriy Kryshtafovych, Torsten Schwede, Maya Topf, Krzysztof Fidelis, and John Moult. Critical assessment of methods of protein structure prediction (CASP)—Round XIV. Proteins: Structure, Function, and Bioinformatics, 89(12):1607–1617, 2021. ISSN 1097-0134. doi: 10.1002/prot.26237. URL https://onlinelibrary.wiley.com/doi/abs/10.1002/prot.26237. eprint: https://onlinelibrary.wiley.com/doi/pdf/10.1002/prot.26237.

[38] Yang Zhang and Jeffrey Skolnick. Scoring function for automated assessment of protein structure template quality. Proteins, 57(4):702–710, December 2004. ISSN 1097-0134. doi: 10.1002/prot.20264.

[39] Sankar Basu and Björn Wallner. DockQ: A Quality Measure for Protein-Protein Docking Models. PLOS ONE, 11(8):e0161879, August 2016. ISSN 1932-6203. doi: 10.1371/journal.pone.0161879. URL https://journals.plos.org/plosone/article?id=10.1371/journal.pone.0161879. Publisher: Public Library of Science.

[40] Roshan Rao, Jason Liu, Robert Verkuil, Joshua Meier, John Canny, Pieter Abbeel, Tom Sercu, and Alexander Rives. MSA Transformer. In Proceedings of the 38th International Conference on Machine Learning, pages 8844–8856. PMLR, July 2021. URL https://proceedings.mlr.press/v139/rao21a.html. ISSN: 2640-3498.

[41] Gustaf Ahdritz, Nazim Bouatta, Sachin Kadyan, Qinghui Xia, William Gerecke, and Mohammed AlQuraishi. OpenFold, November 2021. URL https://zenodo.org/record/6683638.

[42] J. Dauparas, I. Anishchenko, N. Bennett, H. Bai, R. J. Ragotte, L. F. Milles, B. I. M. Wicky, A. Courbet, R. J. de Haas, N. Bethel, P. J. Y. Leung, T. F. Huddy, S. Pellock, D. Tischer, F. Chan, B. Koepnick, H. Nguyen, A. Kang, B. Sankaran, A. K. Bera, N. P. King, and D. Baker. Robust deep learning–based protein sequence design using ProteinMPNN. Science, 378 (6615):49–56, October 2022. doi: 10.1126/science.add2187. URL https://www.science.org/doi/10.1126/science.add2187. Publisher: American Association for the Advancement of Science.

[43] Jue Wang, Sidney Lisanza, David Juergens, Doug Tischer, Joseph L. Watson, Karla M. Castro, Robert Ragotte, Amijai Saragovi, Lukas F. Milles, Minkyung Baek, Ivan Anishchenko, Wei Yang, Derrick R. Hicks, Marc Expòsit, Thomas Schlichthaerle, Jung-Ho Chun, Justas Dauparas, Nathaniel Bennett, Basile I. M. Wicky, Andrew Muenks, Frank Di-Maio, Bruno Correia, Sergey Ovchinnikov, and David Baker. Scaffolding protein functional sites using deep learning. Science, 377(6604):387–394, July 2022. doi: 10.1126/science.abn2100. URL https://www.science.org/doi/abs/10.1126/science.abn2100. Publisher: American Association for the Advancement of Science.

[44] B. I. M. Wicky, L. F. Milles, A. Courbet, R. J. Ragotte, J. Dauparas, E. Kinfu, S. Tipps, R. D. Kibler, M. Baek, F. DiMaio, X. Li, L. Carter, A. Kang, H. Nguyen, A. K. Bera, and D. Baker. Hallucinating protein assemblies, June 2022. URL https://www.biorxiv.org/content/10.1101/2022.06.09.493773v1. Pages: 2022.06.09.493773 Section: New Results.

[45] Richard Evans, Michael O’Neill, Alexander Pritzel, Natasha Antropova, Andrew Senior, Tim Green, Augustin Žídek, Russ Bates, Sam Blackwell, Jason Yim, Olaf Ronneberger, Sebastian Bodenstein, Michal Zielinski, Alex Bridgland, Anna Potapenko, Andrew Cowie, Kathryn Tunyasuvunakool, Rishub Jain, Ellen Clancy, Pushmeet Kohli, John Jumper, and Demis Hassabis. Protein complex prediction with AlphaFold-Multimer, March 2022. URL https://www.biorxiv.org/content/10.1101/2021.10.04.463034v2. Pages: 2021.10.04.463034 Section: New Results.

[46] Michel van Kempen, Stephanie S. Kim, Charlotte Tumescheit, Milot Mirdita, Cameron L.M. Gilchrist, Johannes Söding, and Martin Steinegger. Foldseek: fast and accurate protein structure search. February 2022. doi: 10.1101/2022.02.07.479398. URL http://biorxiv.org/lookup/doi/10.1101/2022.02.07.479398.

[47] Simon C Potter, Aurélien Luciani, Sean R Eddy, Youngmi Park, Rodrigo Lopez, and Robert D Finn. HMMER web server: 2018 update. Nucleic Acids Research, 46(Web Server issue):W200–W204, July 2018. ISSN 0305-1048. doi: 10.1093/nar/gky448. URL https://www.ncbi.nlm.nih.gov/pmc/articles/PMC6030962/.

[48] Martin Steinegger and Johannes Sö ding. MMseqs2 enables sensitive protein sequence searching for the analysis of massive data sets. Nature Biotechnology, 35(11):1026–1028, November 2017. ISSN 1546-1696. doi: 10.1038/nbt.3988. URL https://www.nature.com/articles/nbt.3988. Number: 11 Publisher: Nature Publishing Group.

[49] Tristan Bepler and Bonnie Berger. Learning the protein language: Evolution, structure, and function. Cell Systems, 12(6):654–669.e3, June 2021. ISSN 2405-4712. doi: 10.1016/j.cels.2021.05.017. URL https://www.sciencedirect.com/science/article/pii/S2405471221002039.

[50] Ethan C. Alley, Grigory Khimulya, Surojit Biswas, Mohammed AlQuraishi, and George M. Church. Uni-fied rational protein engineering with sequence-only deep representation learning. Nature Methods, 12: 1315–1322, March 2019. ISSN 15487105. doi: 10.1101/589333. URL https://www.biorxiv.org/content/10.1101/589333v1. Publisher: Cold Spring Harbor Laboratory.

[51] Michael Heinzinger, Ahmed Elnaggar, Yu Wang, Christian Dallago, Dmitrii Nechaev, Florian Matthes, and Burkhard Rost. Modeling the language of life – Deep Learning Protein Sequences. bioRxiv, page 614313, 2019. doi: 10.1101/614313. URL https://www.biorxiv.org/content/10.1101/614313v3. Publisher: Cold Spring Har-bor Laboratory.

[52] Roshan Rao, Nicholas Bhattacharya, Neil Thomas, Yan Duan, Xi Chen, John Canny, Pieter Abbeel, and Yun S. Song. Evaluating Protein Transfer Learning with TAPE. In Neural Information Processing Systems. Cold Spring Harbor Labo-ratory, June 2019. doi: 10.1101/676825. URL https://doi.org/10.1101/676825http://arxiv.org/abs/1906.08230.

[53] Ahmed Elnaggar, Michael Heinzinger, Christian Dallago, Ghalia Rihawi, Yu Wang, Llion Jones, Tom Gibbs, Tamas Feher, Christoph Angerer, Debsindhu Bhowmik, and Burkhard Rost. ProtTrans: Towards Cracking the Language of Lifes Code Through Self-Supervised Deep Learning and High Performance Computing. IEEE Transactions on Pattern Analy-sis and Machine Intelligence, 14(8), July 2021. doi: 10.1109/TPAMI.2021.3095381. URL https://www.osti.gov/pages/biblio/1817585. In-stitution: Oak Ridge National Lab. (ORNL), Oak Ridge, TN (United States).

[54] Joshua Meier, Roshan Rao, Robert Verkuil, Jason Liu, Tom Sercu, and Alexander Rives. Language models enable zero-shot prediction of the effects of mutations on protein function. preprint, Synthetic Biology, July 2021. URL http://biorxiv.org/lookup/doi/10.1101/2021.07.09.450648.

[55] Brian L Hie, Kevin K Yang, and Peter S Kim. Evolu-tionary velocity with protein language models predicts evolutionary dynamics of diverse proteins. Cell Sys-tems, 13(4):274–285.e6, 2022.

[56] Brian Hie, Ellen D. Zhong, Bonnie Berger, and Bryan Bryson. Learning the language of viral evolution and escape. Science, 371(6526):284–288, January 2021. ISSN 1095-9203. doi: 10.1126/science.abd7331.

[57] Karim Beguir, Marcin J. Skwark, Yunguan Fu, Thomas Pierrot, Nicolas Lopez Carranza, Alexan-dre Laterre, Ibtissem Kadri, Abir Korched, Anna U. Lowegard, Bonny Gaby Lui, Bianca Sänger, Yun-peng Liu, Asaf Poran, Alexander Muik, and Ugur Sahin. Early Computational Detection of Potential High Risk SARS-CoV-2 Variants, September 2022. URL https://www.biorxiv.org/content/10.1101/2021.12.24.474095v2.

[58] Nadav Brandes, Grant Goldman, Charlotte H. Wang, Chun Jimmie Ye, and Vasilis Ntranos. Genome-wide prediction of disease variants with a deep protein language model, August 2022. URL https://www.biorxiv.org/content/10.1101/2022.08.25.505311v1.

[59] Erik Nijkamp, Jeffrey Ruffolo, Eli N Weinstein, Nikhil Naik, and Ali Madani. Progen2: exploring the bound-aries of protein language models. arXiv preprint arXiv:2206.13517, 2022.

[60] Mohammed AlQuraishi. End-to-End Differentiable Learning of Protein Structure. (4):292–301.e3, April 2019. doi: 10.1016/j.cels.2019.03.006. Cell Systems, 8 ISSN 2405-4712. URL https://www.sciencedirect.com/science/article/pii/S2405471219300766.

[61] Yang Zhang. Protein structure prediction: when is it useful? Current Opinion in Structural Biology, 19(2):145–155, April 2009. ISSN 1879-033X. doi: 10.1016/j.sbi.2009.02.005.

[62] Chloe Hsu, Robert Verkuil, Jason Liu, Zeming Lin, Brian Hie, Tom Sercu, Adam Lerer, and Alexander Rives. Learning inverse folding from millions of predicted structures. In Proceedings of the 39th International Conference on Machine Learning, pages 8946–8970. PMLR, June 2022. URL https://proceedings.mlr.press/v162/hsu22a.html. ISSN: 2640-3498.

[63] Patrick Kunzmann and Kay Hamacher. Biotite: a unifying open source computational biology framework in Python. BMC Bioinformatics, 19(1):346, October 2018. ISSN 1471-2105. doi: 10.1186/s12859-018-2367-z. URL https://doi.org/10.1186/s12859-018-2367-z.

[64] S. D. Dunn, L. M. Wahl, and G. B. Gloor. Mutual information without the influence of phylogeny or entropy dramatically improves residue contact prediction. Bioinformatics, 24(3):333–340, February 2008. ISSN 13674803. doi: 10.1093/bioinformatics/btm604. Publisher: Oxford Academic.

[65] Fabian Pedregosa, Gaël Varoquaux, Alexandre Gramfort, Vincent Michel, Bertrand Thirion, Olivier Grisel, Mathieu Blondel, Peter Prettenhofer, Ron Weiss, Vincent Dubourg, Jake Vanderplas, Alexandre Passos, David Cournapeau, Matthieu Brucher, Matthieu Perrot, and Édouard Duchesnay. Scikit-learn: Machine Learning in Python. Journal of Machine Learning Research, 12(85):2825–2830, 2011. ISSN 1533-7928. URL http://scikit-learn.sourceforge.net.

[66] Jianlin Su, Yu Lu, Shengfeng Pan, Bo Wen, and Yunfeng Liu. RoFormer: Enhanced Transformer with Rotary Position Embedding, October 2021. URL http://arxiv.org/abs/2104.09864. arXiv:2104.09864 [cs] version: 2.

[67] Yang You, Jing Li, Sashank Reddi, Jonathan Hseu, Sanjiv Kumar, Srinadh Bhojanapalli, Xiaodan Song, James Demmel, Kurt Keutzer, and Cho-Jui Hsieh. Large Batch Optimization for Deep Learning: Training BERT in 76 Minutes. page 38, 2020.

[68] Samyam Rajbhandari, Jeff Rasley, Olatunji Ruwase, and Yuxiong He. ZeRO: Memory optimizations Toward Training Trillion Parameter Models. n SC20: International Conference for High Performance Computing, Networking, Storage and Analysis, pages 1–16, November 2020. doi: 10.1109/SC41405.2020.00024.

[69] Jonathan Ho, Nal Kalchbrenner, Dirk Weissenborn, and Tim Salimans. Axial Attention in Multidimensional Transformers. arXiv, December 2019. URL http://arxiv.org/abs/1912.12180. Publisher: arXiv.

[70] F. Alexander Wolf, Philipp Angerer, and Fabian J. Theis. SCANPY: large-scale single-cell gene expression data analysis. Genome Biology, 19(1):15, February 2018. ISSN 1474-760X. doi: 10.1186/s13059-017-1382-0. URL https://doi.org/10.1186/s13059-017-1382-0.

[71] Isaac Virshup, Sergei Rybakov, Fabian J. Theis, Philipp Angerer, and F. Alexander Wolf. anndata: Annotated data, December 2021. URL https://www.biorxiv.org/content/10.1101/2021.12.16.473007v1. Pages: 2021.12.16.473007 Section: New Results.

[72] Leland McInnes, John Healy, and James Melville. UMAP: Uniform Manifold Approximation and Projection for Dimension Reduction, September 2020. URL http://arxiv.org/abs/1802.03426. arXiv:1802.03426 [cs, stat].

